# Advancing Provenance Assignment using Machine Learning and Time Series Analysis of Chemical Chronologies in Archival Tissues

**DOI:** 10.1101/2025.08.19.671102

**Authors:** Kohma Arai, Malte Willmes, Rachel C. Johnson, Anna M. Sturrock

## Abstract

1. Accurately assigning the provenance of organisms is critical for understanding ecological connectivity and guiding effective conservation and management. Natural chemical chronologies stored in metabolically inert, incrementally growing tissues (e.g., otoliths) provide a powerful tool for this purpose. However, traditional approaches face biological challenges (e.g., dispersal, maternal effects), collapse chronological data into summary metrics, and rely on subjective interpretation—limiting their accuracy and scalability.
2. We present a novel, flexible framework that integrates machine learning, time-series analysis, and ensemble modeling to improve provenance assignment from archival tissue chemistry. Using otolith ^87^Sr/^86^Sr profiles from 17 natal sources of California Central Valley Chinook salmon (*Oncorhynchus tshawytscha*), we moved beyond conventional summary-based methods by developing fully automated time-series feature extraction, explicit time-series classification (including dynamic time warping [DTW] with *k*-nearest neighbors [KNN]), and ensemble models that combine multiple classifiers. We further incorporated simulated data to represent under-sampled life history strategies and validated the framework on real-world, known- and unknown-origin samples.
3. Time-series-based approaches consistently outperformed traditional methods, particularly for sources with strong maternal signatures or early dispersal. Feature extraction approaches informed by biological knowledge were most effective when chemical chronologies followed predictable life-stage-specific patterns, whereas explicit time-series classification (DTW + KNN) excelled when sources displayed distinct overall profile “shapes”. Ensemble models leveraged the complementary strengths of individual approaches, outperforming any single method. Incorporating simulated data corrected systematic underrepresentation of key life history phenotypes in real-world applications, improving model performance, population composition estimates, and their relevance for management decisions.
4. Our results highlight the power of treating archival chemical data as time series, combined with machine learning and ensemble strategies, to enhance the accuracy, consistency, and scalability of provenance assignment. This flexible framework is broadly applicable across taxa, tissues, and chemical markers, offering a practical roadmap for advancing ecological inference and informing conservation and management.

## 1. Introduction

Tracing the origins and movements of organisms is key to understanding ecological processes and informing conservation and management. Provenance tools are widely used in contexts ranging from migration and connectivity studies to wildlife forensics, food traceability, and species management (Bataille et al., 2020; Carter & Chesson, 2017; Hobson, 1999). While population genetics can distinguish long-diverged groups, even modest mixing can obscure genetic structure at management-relevant scales (Christie et al., 2017). A growing alternative uses chemical records in metabolically inert, incrementally growing tissues such as bird feathers (van Dijk et al., 2014), mammal teeth (Newsome et al., 2010), and fish otoliths, eye lenses, vertebrae, and scales (Reis-Santos et al., 2023). In aquatic organisms, the chemistry of these tissues reflects the physicochemical properties of ambient water conditions (Hüssy et al., 2020), enabling reconstruction of provenance and movement histories across taxa and ecosystems (Doubleday et al., 2022). However, robust and flexible analytical frameworks are still needed to translate these chemical signals into reliable provenance estimates.

Two major biological challenges limit the effectiveness of natural tags. First, mobile species may disperse before the relevant structure has recorded the natal chemical signal (Fig. 1). This is especially problematic in marine organisms, where eggs and larvae are passively transported long distances by ocean currents (Cowen & Sponaugle, 2009). Many powerful geolocators in marine waters (e.g., carbonate δ^18^O) require considerable material, often integrating signals over long timescales (Hanson et al., 2010). As a result, many studies focus on broader spatial units such as post-settlement juvenile nursery areas (e.g., Arai et al., 2021). Within-population variation in migration behavior (i.e., partial migration; Chapman et al., 2011) presents an additional challenge. Even when individuals originate from the same natal region, differences in migration strategies (e.g., early vs. late migrant; Fig. 1) can produce distinct chemical chronologies, complicating provenance assignment. Second, maternal effects can confound natal chemical signals. For egg-laying species, the yolk can nourish offspring for extended periods, resulting in “maternal overprinting” of early chemical signatures (Hegg et al., 2019; Fig. 1). This is particularly problematic for species with larger eggs and where maternal environments differ from those of the offspring at the start of life. For example, anadromous salmonids undergo vitellogenesis in the ocean but spawn in freshwater rivers, creating a salinity-based mismatch between the chemical signature of the yolk versus the offspring’s natal environment. This mismatch, combined with early dispersal, can result in chemical chronologies that never reflect the natal signature of the offspring. While useful for inferring maternal strategies (e.g., resident vs. anadromous; Courter et al., 2013) or for artificial marking (Almany et al., 2007), maternal signals can complicate provenance identification.

**Figure 1.**
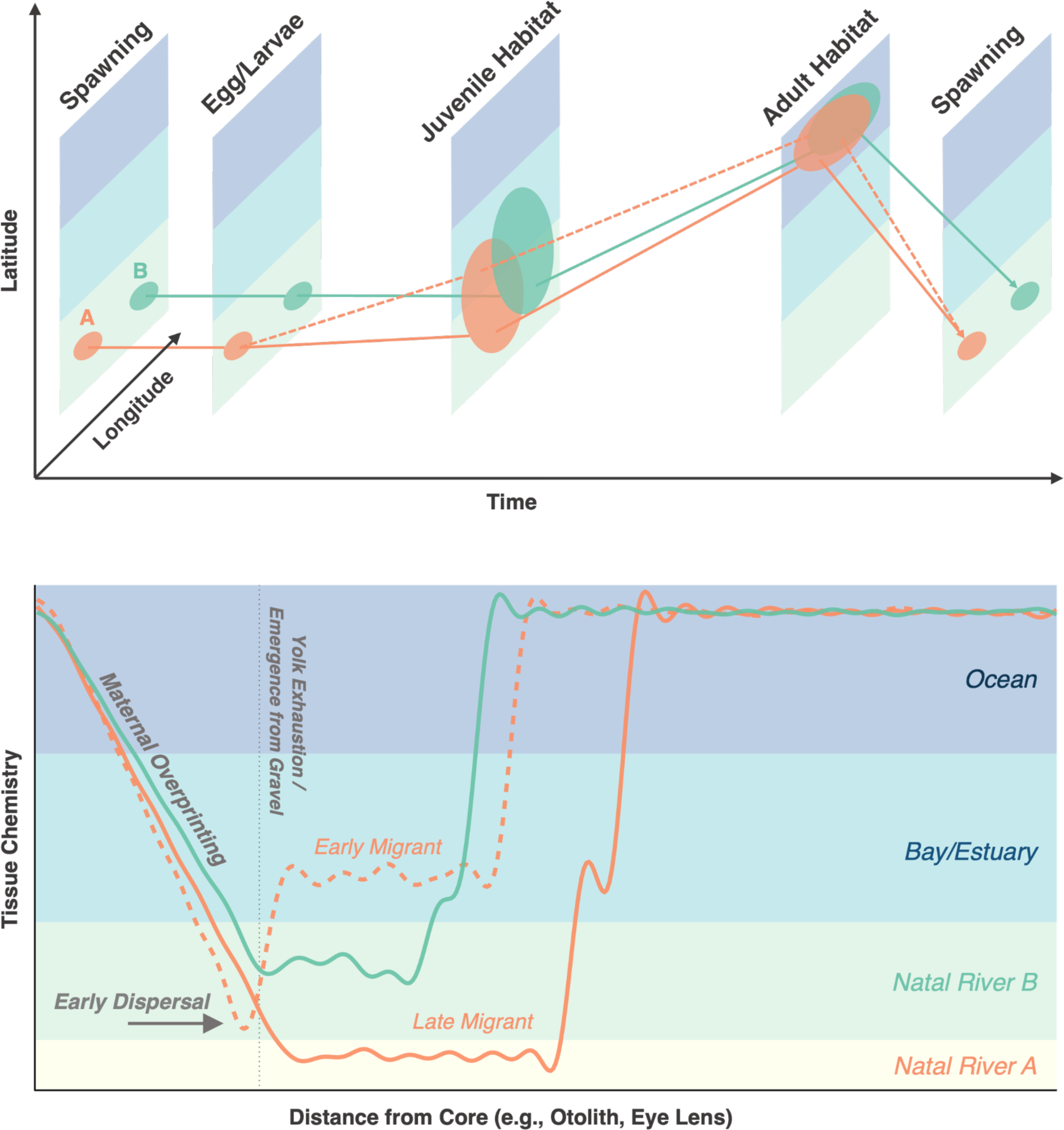
Schematic showing geographic distributions of life history stages and corresponding chemical chronologies in archival tissues (e.g., otolith, eye lens) of anadromous salmonids. Life histories for two distinct natal sources (A and B) are depicted. For source A, fish exhibit early (dashed) or late (solid line) downstream migration. (i) Early migrants may leave the natal river before the tissue records its chemical signature. (ii) The initial portion of the time series reflects maternal chemical signature (“maternal overprinting”) until yolk reserves are exhausted (dotted line), after which ambient signals emerge. Ellipses represent potential geographic distributions; background colors indicate different habitats. Note that this example was inspired by otolith ^87^Sr/^86^Sr (uniform at sea), but for markers exhibiting spatial variation among adult feeding grounds (e.g., δ^13^C), their offspring’s archival structures could reflect such differences in the core. Figure inspired by Figure 1 in Secor (1999).

In addition to these biological constraints, methodological challenges limit the scalability of chemical records for provenance assignment. For instance, visual stress checks in otoliths can provide valuable markers of key life history transitions, such as emergence or settlement, and are commonly used to guide targeted chemical analyses (Barnett-Johnson et al., 2007). However, these checks do not always align with natal signature formation. For example, a larva displaced by high flows may not deplete its yolk and form a stress mark until well after it has left its natal area, potentially decoupling chemical signals from geographic origin. Moreover, identifying regions of interest within chemical time series typically relies on manual inspection (Brennan et al., 2015), introducing subjectivity, limiting throughput, and constraining use in large-scale applications.

A further limitation in the analysis of chemical time series is the tendency to oversimplify their inherently chronological structure. Rather than leveraging the full sequence of measurements, many studies reduce these data to summary statistics (e.g., mean values) within a fixed window, leading to a loss of temporal resolution and biological nuance (Barnett-Johnson et al., 2008). Yet time series contain information not only in the individual values, but also in their sequence and overall structure, all of which are obscured when condensed into static summaries (Chatfield & Xing, 2019).

Time-series methods are increasingly used to identify migratory phenotypes (e.g., Cordoleani et al., 2021), and more recently been applied to characterize larval trajectories and partial migration using unsupervised clustering such as Dynamic Time Warping (DTW; Arai et al., 2024; Hegg & Kennedy, 2021; Teichert et al., 2024). Machine learning classifiers such as random forests (RF) and *k*-nearest neighbors (KNN) outperform traditional methods (e.g., linear discriminant analysis) in capturing non-linear relationships and temporal patterns (Arai et al., 2023; Mercier et al., 2011). Ensemble approaches, which integrate multiple classifiers, are also gaining popularity for their ability to improve predictive accuracy by leveraging the strength of diverse models (Dietterich, 2000).

A key assumption in supervised natal assignment is that all contributing sources to the mixed group are well represented in the training data (Elsdon et al., 2008). However, this assumption can break down when rapid dispersal during early life stages or geographically widespread populations makes it logistically challenging to collect sufficient known-origin samples. For these poorly characterized life history traits or natal sources, the lack of real, direct observations can introduce substantial bias into provenance models. In such cases, simulation techniques based on established biological principles, such as chemical uptake dynamics, known water chemistries, and fish movement patterns, can provide a realistic proxy for actual measurements, effectively serving as a reference dataset for model training (Artetxe-Arrate et al., 2025). Together, these approaches offer a promising framework for overcoming the biological and technical challenges inherent in provenance assignment from chemical time series in archival tissues.

Otolith ^87^Sr/^86^Sr ratios are particularly effective for tracking fish movements among rivers with distinct geologies, as these isotopic ratios vary predictably with underlying bedrock and incorporated reliably into otoliths (Barnett-Johnson et al., 2008; Bataille & Bowen, 2012). California’s Central Valley, with its geologically diverse watersheds, is ideal for such provenance applications (Barnett-Johnson et al., 2008; Johnson et al., 2016). California Central Valley fall-run Chinook salmon (*Oncorhynchus tshawytscha*) display diverse life history traits, both spatially (across distinct spawning populations) and temporally (through varied juvenile emigration timings within populations; Williams, 2006). Otolith ^87^Sr/^86^Sr analysis has been widely used to assign natal origin and estimate juvenile outmigration timing (Sturrock et al., 2020; Willmes et al., 2024). However, juveniles that leave natal rivers shortly after emergence (early migrants) may retain maternal signals and lack sufficient time to incorporate distinct natal river signatures into their otoliths, making them particularly susceptible to misclassification using traditional methods (Sturrock et al., 2020; Willmes et al., 2024; see also Fig. 1). These early migrants are typically too small to tag at the time of dispersal, and if later sampled downstream, their natal origin cannot be confirmed, leading to gaps in known-origin reference data for this rapidly dispersing phenotype within a given population. Additionally, conventional assignment approaches rely on manual inspection of individual ^87^Sr/^86^Sr isotope profiles and data summarization, which can reduce both throughput and assignment accuracy.

Here, we use otolith ^87^Sr/^86^Sr time series data from Central Valley Chinook salmon to develop and compare methods for assigning natal origin across 17 distinct sources. We evaluate (1) manual versus automated natal region selection, (2) summary statistics versus full time series representations, and (3) a range of classification techniques. To advance beyond conventional approaches, we introduce novel methodologies that integrate time-series feature extraction, machine learning classification, and ensemble modeling. We also incorporate simulated training data to represent under-sampled early migrant phenotypes and test their contribution to model performance. Finally, we validate our framework using both known-origin and unknown-origin field samples to assess its accuracy and relevance for management decisions. Our goal is to evaluate trade-offs in accuracy, scalability, and efficiency, and to provide a robust, flexible workflow for provenance assignment using archival tissue chemical chronologies.

## 2. Materials and Methods

### 2.1 Datasets

#### Otolith ^87^Sr/^86^Sr isotope reference data

The classification model included 17 natal sources (12 natural, 5 hatchery) that represent the majority of the Central Valley Chinook salmon stock complex. Model training primarily used otolith ^87^Sr/^86^Sr profiles from fall-run juveniles sampled in natal rivers or hatcheries prior to outmigration (Table S1, Fig. 2). For hatcheries, we also included older fish with confirmed origins via coded wire tags (CWT). Hatchery fish were included in the reference baseline because their otolith ^87^Sr/^86^Sr values can differ from those of natural fish from the same river, due to marine-derived feed in hatcheries (Barnett-Johnson et al., 2008).

**Figure 2.**
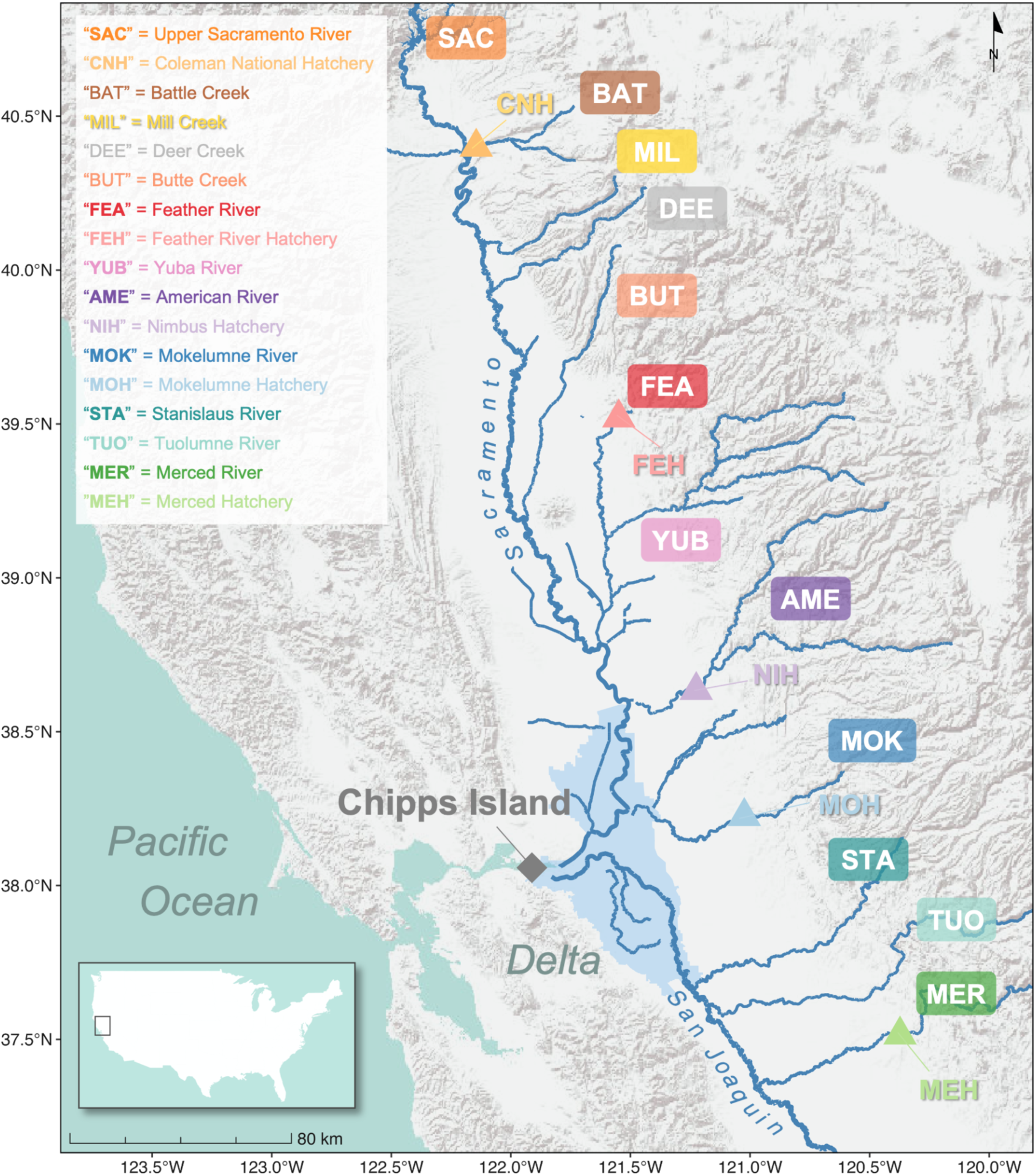
Map of California’s Central Valley showing the locations of salmon-producing rivers and hatcheries (triangles), representing a total of 17 natal sources, along with their corresponding three-letter source codes used throughout the paper. Juvenile Chinook salmon of unknown origin were collected at Chipps Island (diamond), located at the downstream terminus of the Delta.

Due to limited known-origin fall-run juvenile samples for the upper Sacramento River (SAC), Butte (BUT), Mill, (MIL), and Deer (DEE) Creeks, we included otolith ^87^Sr/^86^Sr data from post-spawned adult carcasses recovered during the spawning season—winter-run in the SAC and spring-run in BUT, MIL, and DEE. While not fall-run, these fish likely represent the natal sources of their recovery sites based on high homing rates (Sturrock et al., 2019) and close agreement between their natal otolith signatures and local water chemistry.

Otolith samples from each source were randomly selected across years, resulting in a broad temporal range (Table S1). Temporal variability in ^87^Sr/^86^Sr is expected to be minimal due to the geologic stability of this tracer (Bataille & Bowen, 2012), unlike other elements that may show cohort variation (Gillanders, 2002).

#### Synthetic early migrant data

To address key gaps in the reference dataset, we simulated otolith ^87^Sr/^86^Sr profiles for “early migrants” — juveniles that emigrate shortly after emergence and rear in non-natal habitats. These individuals are underrepresented in empirical data due to their small size and tagging limitations but often dominate outmigration cohorts, making their exclusion a potential source of bias (Sturrock et al., 2020). Simulated profiles were informed by water ^87^Sr/^86^Sr data, otolith growth rates, and migration timing, and incorporated maternal, natal, and downstream phases. We generated five synthetic profiles per natural-origin source (excluding hatcheries, which do not release fish at fry size; Huber & Carlson, 2015). Full simulation procedures are described in Supplementary Text S1 and Table S2.

#### Otolith microstructure features

In addition to otolith ^87^Sr/^86^Sr profiles, we incorporated otolith microstructure features (mean exogenous feeding check score) into the natal assignment model to improve classification accuracy (see Supplementary Text S2 for details).

### 2.2 Approaches for provenance assignment

We developed and compared seven models for assigning salmon natal origin based on otolith ^87^Sr/^86^Sr profile (Fig. 3). These span a continuum from expert-driven, discrete-feature approaches to fully automated, time series-based classification methods. This continuum reflects a progression in how much temporal structure is retained and how much user interpretation is required. The most complex model integrates predictions from multiple approaches into an ensemble, leveraging the strengths of each method.

**Figure 3.**
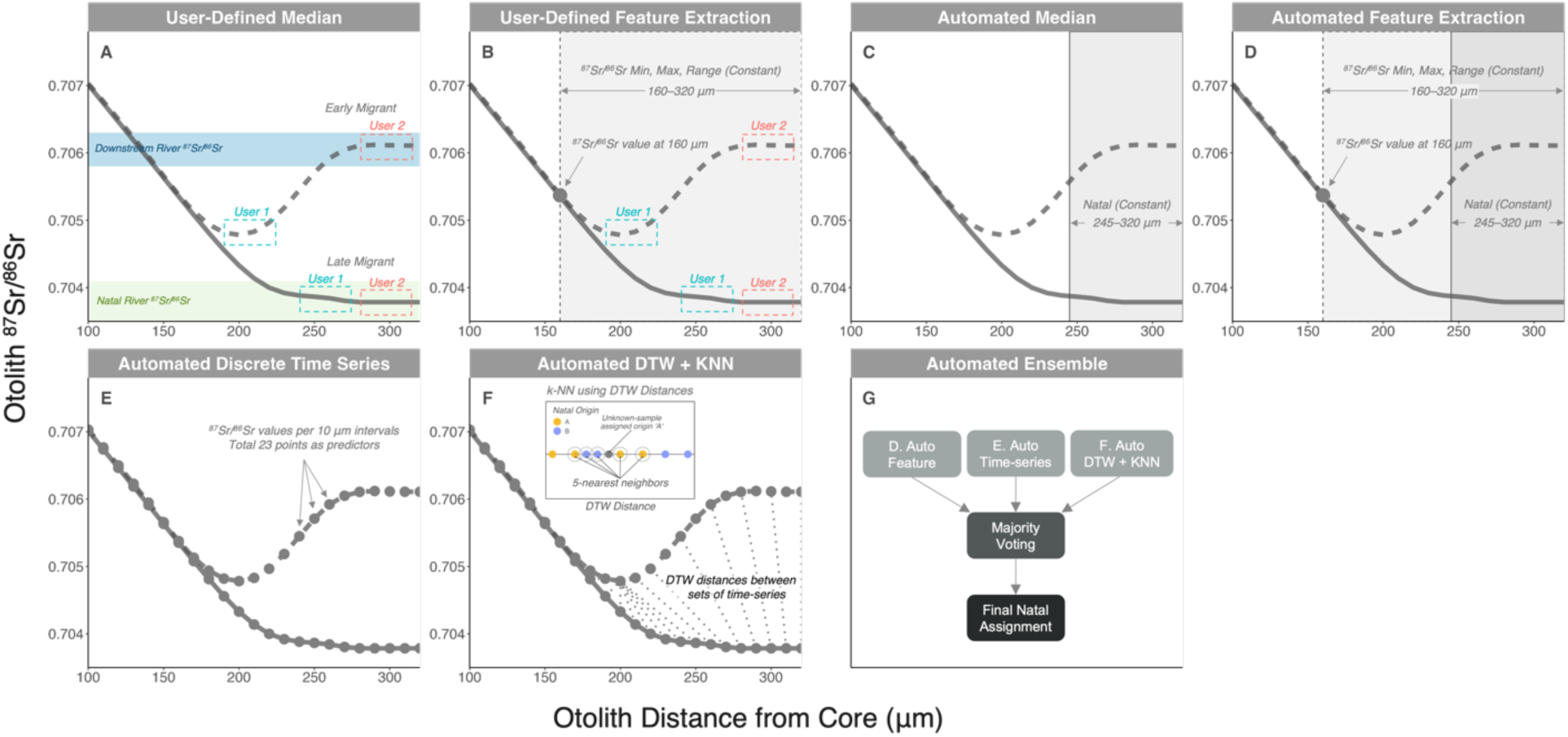
Schematic illustrating the approaches used to assign fish provenance. Solid and dashed lines show otolith ^87^Sr/^86^Sr profiles for late and early migrants, measured from 100 and 320 µm from the core. In panel (A), green and blue shading indicate natal river and downstream river water ^87^Sr/^86^Sr values, respectively. In panels (A) and (B), blue and red dashed boxes indicate user-defined natal regions assigned by two readers. While readers assign similar natal regions for late migrants, they diverge on early migrants, with User 2 selecting a later portion of the profile than User 1. This highlights the subjectivity and inconsistency inherent in conventional manual inspection approaches.

To standardize inputs, we fit spline models to interpolate ^87^Sr/^86^Sr values every 10 µm between 100 and 320 µm from the otolith core. Spline degrees of freedom were adjusted to avoid oversmoothing. We excluded the first 100 µm due to potential overpolishing, which can expose material deposited later in life. We also excluded data beyond 320 µm to avoid the risk of incorporating signatures from non-natal habitats (Sturrock et al., 2020).

#### User-defined models

##### User-defined median

This traditional approach relies on expert visual inspection to identify the natal region—typically the plateau following yolk depletion, often guided by the exogenous feeding check (Fig. 3A). The median ^87^Sr/^86^Sr value within this region, plus the mean exogenous feeding check score, were used as predictors in a random forest (RF) classifier. While flexible, this approach is time-intensive and subjective, as each profile must be interpreted manually.

##### User-defined feature extraction

To account for early downstream movements of fish—before their otoliths fully reflect the natal ^87^Sr/^86^Sr signature—this model builds upon the user-defined median approach by incorporating additional time series features derived from prior biological knowledge of Chinook salmon in this system (Fig. 3B). Although this feature extraction approach is not a time series method in the formal sense, it attempts to extract biologically meaningful time series features informed by expert knowledge. Specifically, we calculated the minimum, maximum, and range of ^87^Sr/^86^Sr values between 160 and 320 µm, based on the premise that these features could capture movements across habitats with distinct ^87^Sr/^86^Sr signatures unique to each source. We also included the ^87^Sr/^86^Sr value measured at 160 µm from the core, as this point often precedes the onset of early dispersal, and thus all individuals from a given source tend to share similar isotopic values at this size. These summary statistics were used alongside the user-defined median ^87^Sr/^86^Sr and the mean exogenous feeding check score in RF classification.

##### Automated discrete models

To reduce subjectivity and increase efficiency, we developed automated variants of the above two models using fixed rules to define the natal region and extract summary statistics.

##### Automated median

This model defines a fixed natal region (245320 µm) for all samples assumed to represent otolith growth post-yolk absorption but prior to outmigration (Fig. 3C; Barnett-Johnson et al., 2008). The median ^87^Sr/^86^Sr value within this region and the mean exogenous feeding check score were used as RF inputs.

##### Automated feature extraction

This model mirrors the automated median approach but includes additional features to capture variability in the ^87^Sr/^86^Sr profile (Fig. 3D). In addition to the median ^87^Sr/^86^Sr value and mean exogenous feeding check score, we calculated the minimum, maximum, and range of ^87^Sr/^86^Sr values between 160 and 320 µm, as well as the ^87^Sr/^86^Sr value measured at 160 µm from the core and used as RF inputs.

#### Time series models

Moving beyond summary statistics, the next set of models retain the temporal structure of the otolith ^87^Sr/^86^Sr profiles.

##### Discrete time-series

This model treats the otolith ^87^Sr/^86^Sr profile as a set of interpolated values sampled at discrete intervals (Fig. 3E). Specifically, 23 ^87^Sr/^86^Sr values spaced every 10 µm from 100 to 320 µm were used as individual predictors in RF, alongside the mean exogenous feeding check score. Although this automated approach does not require a user-defined natal region and leverages regularly spaced values from a time series, it does not explicitly model the sequential nature of the data.

##### Dynamic time warping with *k*-nearest neighbor classification (DTW + KNN)

To explicitly treat the otolith ^87^Sr/^86^Sr profile as a continuous time series and to classify samples based on the overall structure and sequence of the profile, we developed a model that combines dynamic time warping (DTW) with *k*-nearest neighbor (KNN) classification (Wang et al., 2013; Fig. 3F). DTW aligns sequences by minimizing time-warping cost and enables comparison of profiles with similar patterns but varying lengths (Aghabozorgi et al., 2015; Hegg & Kennedy, 2021). While DTW has been applied to otolith chemistry data in an unsupervised context to identify distinct migration patterns (Arai et al., 2024; Hegg et al., 2019; Teichert et al., 2024), our approach represents a novel application of DTW for supervised classification by pairing it with KNN. This allows us to directly classify unknown samples based on their temporal structure, leveraging the full profile information rather than relying on summary statistics or discrete points.

We computed the DTW distance between the ^87^Sr/^86^Sr profile of each unknown-origin sample and all known-origin samples in the training set. Classification was then performed using KNN (*k* = 5; see below), where the sample was assigned to the most common class among its five nearest neighbors based on DTW distance. Note that this method did not incorporate information about the mean exogenous feeding check score, and assignments were based solely on DTW distance.

#### Ensemble model

Finally, to capitalize on the complementary strengths of different modeling strategies, we developed an ensemble classifier that integrates predictions from three automated approaches: the automated feature extraction, discrete time-series, and DTW + KNN models (Fig. 3G). Each method independently generated a class prediction, and the final assigned natal origin was determined by majority vote among the three predictions.

We included a “null” model that assigns all samples to the most frequent class to benchmark the classification performance of all other classifiers. Natal assignments were performed in R (R Core Team, 2025) using *tidymodels* (Kuhn & Wickham, 2020), *ranger* (Wright & Ziegler, 2017), *dtw* (Giorgino, 2009), *fastKNN* (Besanson, 2015), and *manymodelsR* (Gonzabato, 2025). Hyperparameters for RF *(mtry* and *min_n*) and KNN (*k*) were tuned using grid search with 10-fold cross-validation. The full set of optimal hyperparameter combinations is provided in Table S3. Classification accuracy for all approaches was evaluated using 10-fold cross-validation (repeated 10 times).

To provide additional context for model performance, we also compared classification accuracy between the RF and traditional statistical classifiers such as linear and quadratic discriminant analysis, using the automated median approach as a common baseline (Fig. 3C). Detailed methods and results are presented in Supplementary Text S3.

Lastly, to evaluate subjectivity of natal region selection in user-defined approaches, we conducted a visual identification trial in which three trained readers independently selected natal regions based on otolith ^87^Sr/^86^Sr profiles. Detailed methods and results are presented in Supplementary Text S4.

### 2.3 Natal assignment of unknown-origin juveniles

To evaluate the performance of the natal assignment model in a more realistic context, we assigned natal origins to wild, randomly selected unknown-origin juveniles (n = 50) collected at Chipps Island from 2014 to 2021, the downstream terminus of the Delta through which all natal river sources must pass to reach the ocean (Fig. 2; Table S1). These fish were collected under scientific permit SC-12811.

### 2.4 Natal assignment of known-origin adult returns

To assess the potential benefit of including simulated synthetic early migrants in the training data, we tested model performance using an independent set of otolith ^87^Sr^/86^Sr profile data from post-spawned adults sampled from the Yuba (YUB), American (AME), and Stanislaus (STA) Rivers (n = 30; Fig. 2; Table S1). These adults were assumed to be known-origin, early migrants based on high homing rates and chemical transitions between natal rivers and subsequent habitats along their migration path. We specifically compared the classification accuracy of models trained with and without synthetic early migrants to assess the impact of this additional training data.

## 3. Results

### 3.1 Otolith ^87^Sr/^86^Sr of Central Valley Chinook salmon

The California Central Valley’s geologically diverse watershed produced unique otolith ^87^Sr/^86^Sr profiles for Chinook salmon from 17 natal sources (Fig. 4, Fig. S1). Both natural-origin and hatchery fish showed ^87^Sr/^86^Sr values close to maternal marine values near the otolith core, gradually transitioning to natal river signatures as yolk reserves were exhausted. The steepness of this transition reflected the degree of difference between ocean and natal water chemistry: sources with highly distinct natal signatures (e.g., BAT) showed steep transitions, while those closer to marine values (e.g., MEH) exhibited more gradual shifts.

**Figure 4.**
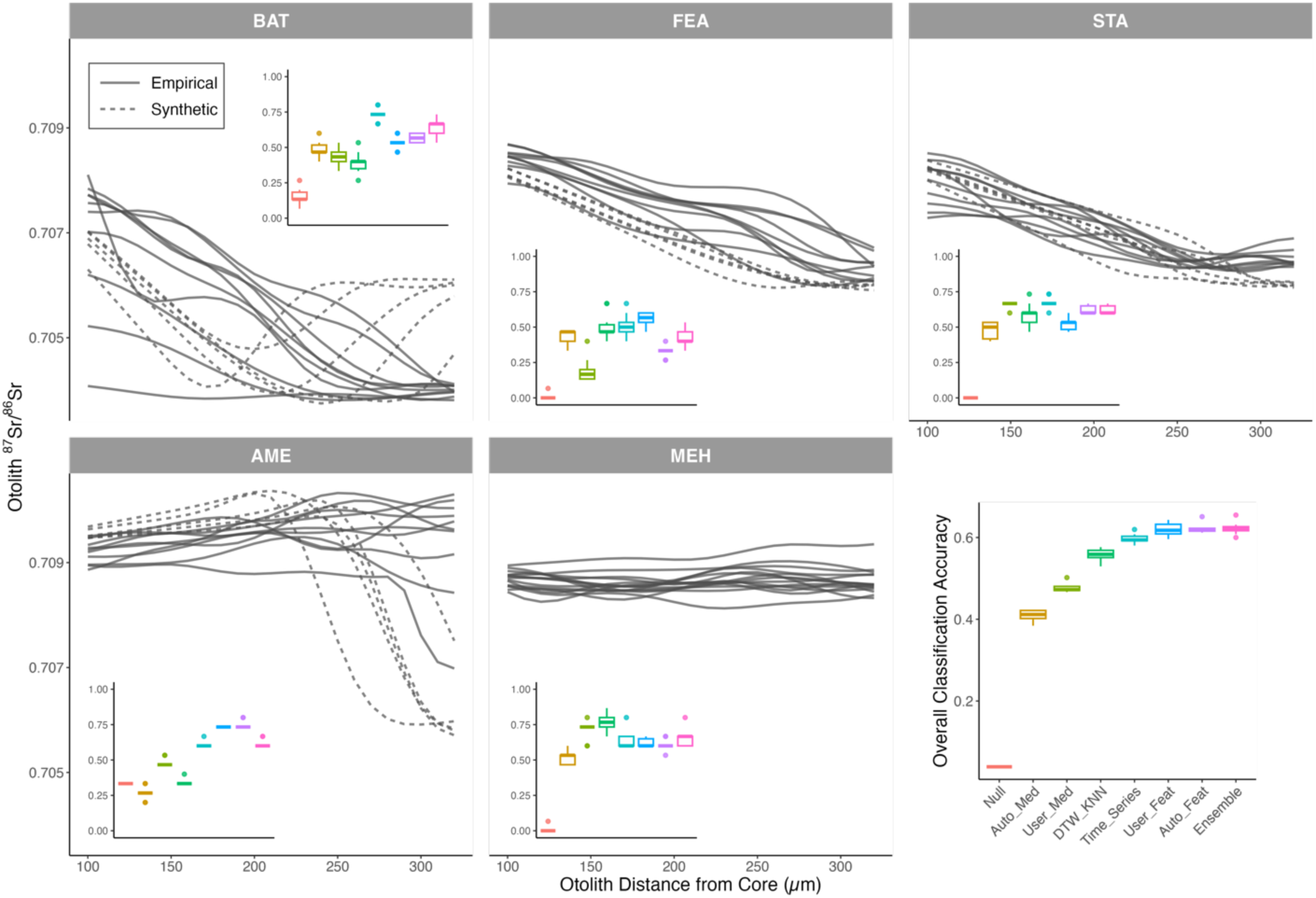
Classification accuracy of natal assignment models evaluated using repeated 10-fold cross-validation (n = 255 known-origin Chinook salmon). Inset plots show otolith ^87^Sr/^86^Sr profiles by natal source, with classification accuracy indicated for each. Solid and dashed lines represent empirical and synthetic fish, respectively. Overall classification accuracy is shown in the bottom right panel. This figure displays a subset of natal sources (5 out of 17) chosen to illustrate a range of distinct otolith ^87^Sr/^86^Sr patterns; results for all 17 sources, including classification accuracy and otolith ^87^Sr/^86^Sr profiles, are shown in Figure S1. Model abbreviations: “Null” = non-informative model; “Auto_Med” / “User_Med” = automated / user-defined median; “DTW_KNN” = dynamic time warping with *k*-nearest neighbors; “Time_Series” = discrete time series; “User_Feat” / “Auto_Feat” = user-defined / automated feature extraction; “Ensemble” = ensemble model. Natal source abbreviations are defined in Figure 2.

Differences in otolith ^87^Sr/^86^Sr profiles between early and late migrants also varied by source. In rivers with natal signatures distinct from the downstream Delta, early migrants exhibited pronounced shifts compared to late migrants (e.g., AME). In contrast, sources with values similar to the Delta showed minimal differences (e.g., FEA).

### 3.2 Classification accuracy among different approaches

Ten-fold cross-validation, repeated 10 times on known-origin juvenile samples, showed that the ensemble (mean ± SD: 0.622 ± 0.015), automated feature extraction (0.622 ± 0.011), and user-defined feature extraction (0.619 ± 0.016) models had the highest classification accuracy (Fig. 4). All models substantially outperformed the “null” model (0.039), indicating they captured meaningful structure in the data beyond random chance.

Accuracy varied by natal source, with each model showing unique strengths (Fig. 4, Fig. S1). The median-based approaches were generally outperformed, but performed moderately well for hatchery sources with distinct ^87^Sr/^86^Sr values between 245–320 µm from the otolith core (e.g., MEH), or where early and late migrants had similar ^87^Sr/^86^Sr profiles (e.g., STA). The automated feature extraction performed well across most sources, particularly where early migrants differed markedly from late migrants within the same source (e.g., AME). The discrete time-series and DTW + KNN approaches excelled where profiles lacked distinct values at fixed points but had unique overall “profile shapes” (e.g., FEA, MEH). The ensemble model, which combined predictions from the automated feature extraction, discrete time-series, and DTW + KNN approaches, consistently performed well across sources.

### 3.3 Natal assignment of unknown-origin juveniles

Applying different natal assignment models to unknown-origin juveniles collected at Chipps Island (the Delta terminus where all juvenile sources pass to reach the ocean; Fig. 2) from 2014 to 2021 revealed a diverse array of sources contributing to the stock complex (Fig. 5; Fig. S2). Among-model assignment agreement varied from 0.34 to 0.92 (Fig. S3). The DTW + KNN, discrete time series, user-defined feature extraction, automated feature extraction, and ensemble approaches generally produced consistent assignments (agreement: 0.64–0.92). In contrast, the lowest agreement (0.34–0.52) occurred between median-based models (e.g., automated or user-defined median) and time-series based approaches (e.g., discrete time series and DTW + KNN).

**Figure 5.**
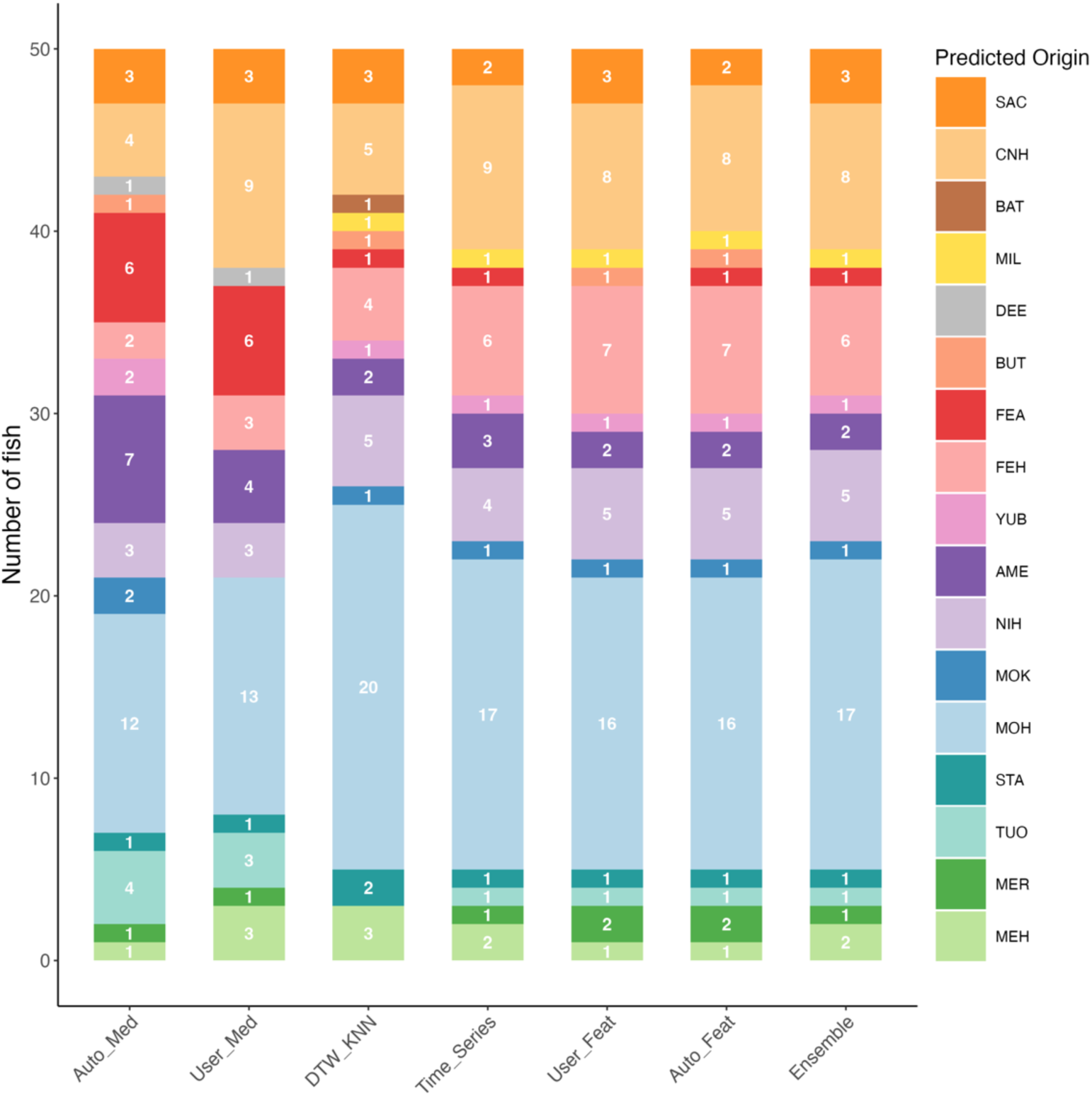
Natal assignment results using different modeling approaches, applied to unknown-origin juvenile Chinook salmon (n = 50) collected at Chipps Island. The numbers in each bar represent the sample size assigned to each predicted natal source. These samples were randomly selected from fish collected between 2014 and 2021 and do not reflect the true population composition at Chipps Island in any specific year, as the primary purpose was to evaluate model performance with real-world samples. Model abbreviations are defined in Figure 4.

### 3.4 Natal assignment of known-origin adult returns

The user-defined feature extraction approach yielded the highest overall classification accuracy for known-origin adult returns that emigrated early (“early migrants”), performing well both with (accuracy = 0.73) and without (0.57) synthetic early migrants in the training data (Fig. 6; Fig. S4). Accuracy varied by natal sources, with AME showing the highest (0.68), and YUB (0.22) and STA (0.30) the lowest.

**Figure 6.**
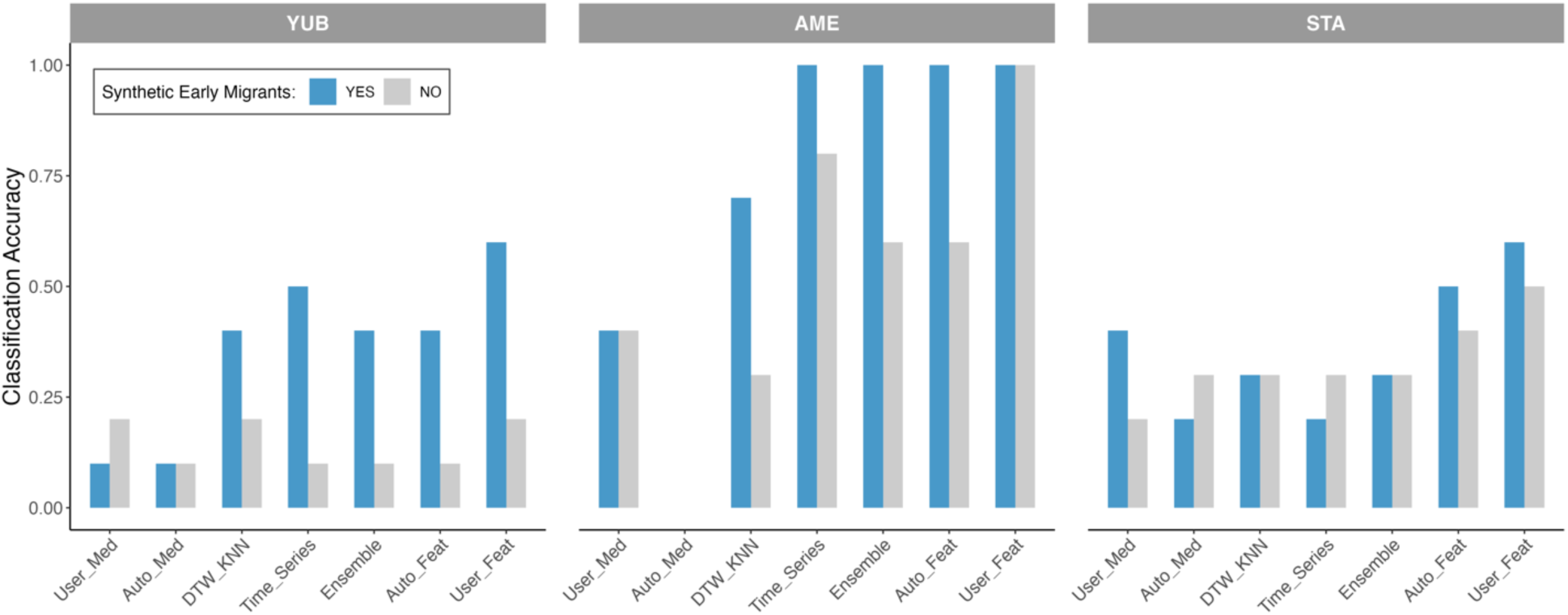
Classification accuracy of known-origin adult returns that emigrated early as fry from the Yuba (YUB), American (AME), and Stanislaus (STA) Rivers, evaluated using different natal assignment models. Results compare models trained with and without synthetic early migrants in the training dataset. Model abbreviations are defined in Figure 4.

Removing synthetic early migrants generally reduced accuracy across models, with the largest declines in the automated feature extraction (−42.1%) and DTW + KNN (−42.9%) models. The effect varied by natal source: accuracy for YUB and AME declined sharply—up to 80% for YUB in the discrete time series approach—while STA showed little to no change. Notably, the user-defined feature extraction model remained robust for AME and STA, showing almost no loss in accuracy after removing synthetic early migrants.

## 4. Discussion

### 4.1 Advancing archival tissue provenance assignment with machine learning and time series approaches

By integrating time-series methods with machine learning, our approach addresses key biological and technical challenges in provenance assignment. Unlike traditional methods that rely on static or single-point measurements, treating chemical data as a time series allowed us to capture temporal structure, improving classification accuracy.

Models that incorporated time series information, either through the discrete time series approach or by including time series-derived features (i.e., user-defined or automated feature extraction approaches), were especially effective where ^87^Sr/^86^Sr profiles exhibited steep transitions from marine to natal values, resulting in larger maternal overprinting (e.g., BAT, SAC; Fig. 4, Fig. S1). Even when profiles did not explicitly reflect natal signatures due to early dispersal, these approaches leveraged the shape and steepness of the initial transition to accurately assign origin, outperforming traditional methods based solely on summary statistics (median-based approaches). Likewise, for sources exhibiting large differences between the natal stream and downstream habitats, resulting in steep transitions in the later part of the profile for early migrants (e.g., BAT, AME), the automated feature extraction model better captured within-source variation than either traditional median-based models or fully explicit time series approaches via DTW + KNN. This demonstrates the advantage of time-series approaches in accounting for early dispersal and maternal influences in natal origin assignment.

While differences in classification accuracy among models were modest, the feature extraction approaches consistently achieved high performance (Fig. 4). These models incorporate time-series structure by extracting summary features (median, minimum, maximum, and range of ^87^Sr/^86^Sr values) rather than preserving the full temporal sequence. By reducing dimensionality, these feature-extraction methods filter out less informative segments of the signal, effectively enhancing the signal-to-noise ratio. Unlike explicit time series approaches—which requires temporal alignment of key chronological features (e.g., yolk depletion or downstream movement) across individuals to maximize assignment accuracy—the feature extraction approach accommodates greater individual variability in the timing of chemical changes. Additionally, the feature extraction approach implicitly incorporates expert knowledge by focusing on specific regions of the profile (e.g., 245–320 µm) where Chinook salmon natal signatures are most likely to be found.

Time-series feature extraction may be especially suitable for species where environmental signals are overlaid onto consistent ontogenetic patterns, a phenomenon commonly observed in otolith chemical records linked to individual physiology (Sturrock et al., 2015). It also suits systems with predictable environmental transitions (e.g., seasonal flows, hypoxia, pollution) that produce characteristic “motifs” in chemical profiles. For example, this approach can be valuable in studies of transboundary migratory species, where chemical records reflect transitions across environmental gradients that occur at consistent developmental stages (Pérez & Hobson, 2007; Walther & Limburg, 2012). The feature extraction approach may also be relevant in studies of partial migration, where individuals from the same source diverge in movement strategy or timing (Chapman et al., 2011). In such cases, extracting features from targeted segments of the time series may outperform analyses using the full profile.

This feature-extraction strategy pairs well with machine learning classifiers such as random forests (RF), which handle large predictor sets, tolerate collinearity, and capture nonlinear relationships more effectively than traditional methods like linear (LDA) or quadratic (QDA) discriminant function analysis (Mercier et al., 2011). In our dataset, RF outperformed LDA and QDA and was more adaptable to complex inputs, which limited the application of traditional statistical models (Supplementary Text S3). Machine learning models offer the flexibility to incorporate multiple features and capture complex, nonlinear interactions. This includes not only multiple chemical measurements from a single tissue but also chemical data from multiple tissues within the same individual (e.g., otoliths, eye lenses, muscle tissue; Rosinski et al., 2023; Walther & Torrance, 2024). These models can also integrate complementary data beyond tissue chemistry, such as otolith microstructure and early growth patterns (Barnett-Johnson et al., 2007), which we included in our classification models, as well as genetic signatures (Brophy et al., 2020).

Explicit time series classification methods, such as DTW + KNN, was particularly effective in distinguishing sources with distinct overall profile “shapes” rather than point-specific markers. This makes DTW + KNN especially powerful in systems with unconstrained chemical baselines and variable movement timing, often shaped by dynamic environmental conditions (Teichert et al., 2024). For example, marine species inhabiting oceanographically dynamic environments may experience highly individualistic exposure histories to changes in temperature, salinity, or prey availability (Secor, 2015). In such cases, methods that retain the integrity of the entire temporal profile—rather than summarizing it through extracted features—can better capture ecologically meaningful variation. Additionally, DTW can readily handle multivariate time series (Aghabozorgi et al., 2015; Arai et al., 2024), making it well suited for cases where multi-element/isotope data are available for each individual, especially in systems where no single marker reliably reflects provenance. Although DTW can be more sensitive to noise and misalignments, they may outperform traditional feature-extraction methods when group differences manifest as patterns rather than as consistent absolute values.

Our findings highlight the advantages of ensemble methods. By combining relatively uncorrelated models (automated feature extraction, discrete time-series, DTW + KNN), the ensemble model consistently outperformed individual approaches. This reflects a key advantage of ensemble methods: when component classifiers are relatively independent, as in our case, aggregating their predictions can reduce variance and improve generalization (Dietterich, 2000). Ensemble methods are especially attractive for their resistance to overfitting and ability to maintain reliable performance across varying conditions. This makes them well-suited for real-world applications like tissue chemical chronologies, where reference datasets may evolve over time. By aggregating diverse classifiers, ensemble models provide a robust solution to temporal shifts in environmental baselines or analytical protocols, helping ensure consistent performance as data structures change.

A major advantage of our framework is the automation and consistency it offers. Manual profile inspection is time-intensive and subjective—our trials revealed up to 40% disagreement in natal assignment among analysts (Supplementary Text 4), underscoring the potential for inconsistent outcomes that could compromise management decisions. In contrast, automated approaches delivered consistent, reproducible assignments with equal or better accuracy. This standardization facilitates scaling up to larger datasets or long-term monitoring programs where manual approaches become impractical.

### 4.2 Real-world performance and management implications

Accurate provenance estimates are essential for spatial management, conservation planning, and food traceability across taxa—including birds, mammals, and fishes (Bataille et al., 2020; Coelho et al., 2017; Hobson, 1999; Johnson et al., 2016). This is particularly critical for Central Valley Chinook salmon, which exhibit diverse life history strategies—spatially distinct spawning regions and variable juvenile migration timing—that buffer the populations against perturbations through the “portfolio effect” (Schindler et al., 2010). Reliable natal origin and population composition estimates are therefore key to understanding how this diversity supports persistence of this highly impacted stock under stressors like drought and altered flows (Sturrock et al., 2020).

Applying a suite of assignment models to unknown-origin juveniles revealed substantial differences in population composition estimates among methods (Fig. 5; Fig. S2). Traditional approaches, like the user-defined median, tended to assign hatchery-origin MOH samples to a natural source (FEA), reflecting its poor classification accuracy for MOH compared to time series-based methods. Such misclassifications could bias management by underestimating hatchery contributions and overestimating natural sources, potentially misguiding restoration priorities, hatchery release strategies, and viability assessments (Johnson et al., 2012).

Analysis of known-origin adult returns further highlight the value of including synthetic early migrants in the training data (Fig. 6). Simulated data especially benefited automated approaches—such as the automated feature extraction and DTW + KNN—showing substantial improvements in classification accuracy. In contrast, user-defined approaches benefited less, likely because expert readers can visually interpret chemical profiles and accurately assign natal origin, even when specific patterns are missing from the empirical reference. Thus, synthetic data designed to capture underrepresented life history variation in empirical samples are especially valuable when expert curation is impractical, or when scaling to larger, more complex datasets.

Incorporating synthetic early migrants also improved population composition estimates (Fig. S4). Excluding them from the training set led to the underrepresentation of early-migrant sources (e.g., AME) and the overrepresentation of others (e.g., FEA) that overlap isotopically with Delta habitats used by early migrants. This mismatch suggests that critical contributors, such as early migrants from AME, may be overlooked if reference datasets fail to capture the full spectrum of life-history strategies. Such omissions could have important consequences for habitat restoration efforts, as they risk excluding ecologically meaningful but underrepresented contributors to the population. In this context, generating synthetic data based on well-established biological principles and local ecological knowledge can be essential for building more complete baselines (but see Neubauer et al., 2013 for a Bayesian alternative).

## 5. Conclusion

By integrating machine learning, time-series classification, and ensemble modeling, we developed a novel framework for provenance assignment that advances beyond traditional expert interpretation and static summary metrics. Our combination of time series feature extraction, dynamic time warping with supervised classification, ensemble approaches, and synthetic training data represents a significant step forward in applying archival tissue chemistry to ecological questions.

Although no single method performed best across all natal sources, time-series feature extraction and ensemble approaches consistently showed strong, robust performance. Optimal method choice will depend on study goals, data structure, and biological context. Together, our findings provide a practical roadmap for selecting and applying provenance assignment strategies and demonstrate the potential for machine learning and time-series tools to advance ecological inference from chemical chronologies.

## Acknowledgements

We thank the many individuals and organizations who contributed samples, data, or support for this study.

*Reference juvenile and adult otoliths and associated chemical data*

We are grateful to Flora Cordoleani (UC Santa Cruz), Carson Jeffres (UC Davis), George Whitman (UC Davis), Justin Glessner (UC Davis Interdisciplinary Center for Plasma Mass Spectrometry), Pedro Morais (UC Berkeley); the California Department of Fish and Wildlife (CDFW; Rob Titus, Tim Heyne, Doug Killam, Clint Garman, Tracy McReynolds, Grantton Henley, Matt Johnson, Ryan Revnak); Pacific States Marine Fisheries Commission (PSMFC); US Fish and Wildlife Service (USFWS; Laurie Earley, Kevin Niemela, Kevin Offill); California Department of Water Resources (DWR; Virginia Afentoulis, Ryon Kurth, Jason Kindopp); Merced Fish Facility (Mary Serr); East Bay Municipal Utility District (EBMUD; Michelle Workman, Ed Ribble); Nimbus Hatchery (Jason Julienne, Gary Novak); FishBio (Jason Guignard, Andrea Fuller); Natalie Stauffer-Olsen (Trout Unlimited); and Cramer Fish Sciences (Joe Merz, Kirsten Sellheim).

*Chipps Island juvenile fish*

We thank the Interagency Ecological Program Delta Juvenile Fish Monitoring Program—USFWS, CDFW, and DWR—for collecting these samples (Jack Ingram, Cory Graham, Jeff McClain, Denise Barnard, Pat Brandes).

*Adult otoliths from the Yuba, Stanislaus, and American rivers*

For the American River analyses, we thank Cramer Fish Sciences (Joe Merz, Kirsten Sellheim, Jamie Sweeney) and the Sacramento Water Forum. For the Stanislaus River analyses, we acknowledge CDFW (Tim Heyne, Steve Tsao, Crystal Sinclair, Gretchen Murphey, Shelly Schubert, Dom Giudice) and the Central Valley Project Improvement Act and USFWS (contract no. 81332-08-G017 to RCJ) under US Bureau of Reclamation Agreement R09AC20043. For the Yuba River analyses, we thank PSMFC (Duane Massa, Colin Laubach) and the Yuba River Management Team.

*Otolith extraction, preparation, and analysis*

We appreciate the many people who supported otolith extraction, preparation, and analysis, including Keiko Mertz, Mollie Ogaz, Laura Coleman, Kelly Neal, Dana Myers, Sierra Schluep, and Bradyn O’Connor.

*Funding and additional contributions*

Funding for analyses and salaries was provided by the CDFW Proposition 1 Grant (Ecosystem Restoration Program and the Water Quality, Supply, and Infrastructure Improvement Act of 2014, CWC §79707[g]) and a UK Research and Innovation Future Leaders Fellowship (MR/V023578/1) to AMS. We also thank Jonathan Walter for providing insightful comments on the initial draft, which greatly improved the quality of this paper.

## Author Contributions

All authors conceived the ideas and designed methodology; Malte Willmes, Rachel Johnson, and Anna Sturrock collected the data; Kohma Arai, Malte Willmes, and Anna Sturrock analysed the data; Kohma Arai and Anna Sturrock led the writing of the manuscript. All authors contributed critically to the drafts and gave final approval for publication.

Statement on inclusion: Our study includes authors from diverse backgrounds, including scientists based in the country where the research was conducted. All authors were involved from the early stages of study design, ensuring that their varied perspectives were integrated from the outset.

## Data Availability

Data and code will be available from the Dryad Digital Repository upon acceptance.

## References

Aghabozorgi, S., Seyed Shirkhorshidi, A., & Ying Wah, T. (2015). Time-series clustering – a decade review. Information Systems, 53, 16–38. 10.1016/j.is.2015.04.007

Almany, G. R., Berumen, M. L., Thorrold, S. R., Planes, S., & Jones, G. P. (2007). Local replenishment of coral reef fish populations in a marine reserve. Science, 316(5825), 742–744. 10.1126/science.1140597

Arai, K., Best, J., Craig, C., Lyubchich, V., Miller, N., & Secor, D. (2024). Early growth and environmental conditions control partial migration of an estuarine-dependent fish. Marine Ecology Progress Series, 732, 149–166. 10.3354/meps14551

Arai, K., Castonguay, M., Lyubchich, V., & Secor, D. H. (2023). Integrating machine learning with otolith isoscapes: Reconstructing connectivity of a marine fish over four decades. PLoS ONE, 18(5), e0285702. 10.1371/journal.pone.0285702

Arai, K., Castonguay, M., & Secor, D. H. (2021). Multi-decadal trends in contingent mixing of Atlantic mackerel (*Scomber scombrus*) in the Northwest Atlantic from otolith stable isotopes. Scientific Reports, 11, 6667. 10.1038/s41598-021-86116-2

Artetxe-Arrate, I., Fraile, I., Lastra-Luque, P., Farley, J., Clear, N., Shahid, U., Razzaque, S. A., Ahusan, M., Vidot, A., Parker, D., Marsac, F., Murua, H., Merino, G., & Zudaire, I. (2025). Otolith stable isotopes highlight the importance of local nursery areas as the origin of recruits to yellowfin tuna (*Thunnus albacares*) fisheries in the western Indian Ocean. Fisheries Research, 281, 107241. 10.1016/j.fishres.2024.107241

Barnett-Johnson, R., Grimes, C. B., Royer, C. F., & Donohoe, C. J. (2007). Identifying the contribution of wild and hatchery Chinook salmon (*Oncorhynchus tshawytscha*) to the ocean fishery using otolith microstructure as natural tags. Canadian Journal of Fisheries and Aquatic Sciences, 64(12), 1683–1692. 10.1139/f07-129

Barnett-Johnson, R., Pearson, T. E., Ramos, F. C., Grimes, C. B., & Bruce MacFarlane, R. (2008). Tracking natal origins of salmon using isotopes, otoliths, and landscape geology. Limnology and Oceanography, 53(4), 1633–1642. 10.4319/lo.2008.53.4.1633

Bataille, C. P., & Bowen, G. J. (2012). Mapping ^87^Sr/^86^Sr variations in bedrock and water for large scale provenance studies. Chemical Geology, 304–305, 39–52. 10.1016/j.chemgeo.2012.01.028

Bataille, C. P., Crowley, B. E., Wooller, M. J., & Bowen, G. J. (2020). Advances in global bioavailable strontium isoscapes. Palaeogeography, Palaeoclimatology, Palaeoecology, 555, 109849. 10.1016/j.palaeo.2020.109849

Besanson, G. (2015). FastKNN: fast k-nearest neighbors [Computer software]. https://CRAN.R-project.org/package=FastKNN

Brennan, S. R., Zimmerman, C. E., Fernandez, D. P., Cerling, T. E., McPhee, M. V., & Wooller, M. J. (2015). Strontium isotopes delineate fine-scale natal origins and migration histories of Pacific salmon. Science Advances, 1(4), e1400124. 10.1126/sciadv.1400124

Brophy, D., Rodríguez-Ezpeleta, N., Fraile, I., & Arrizabalaga, H. (2020). Combining genetic markers with stable isotopes in otoliths reveals complexity in the stock structure of Atlantic bluefin tuna (*Thunnus thynnus*). Scientific Reports, 10(1), 14675. 10.1038/s41598-020-71355-6

Carter, J. F., & Chesson, L. A. (2017). Food forensics: Stable Isotopes as a guide to authenticity and origin. CRC Press.

Chapman, B. B., Brönmark, C., Nilsson, J.-Å., & Hansson, L.-A. (2011). The ecology and evolution of partial migration. Oikos, 120(12), 1764–1775. 10.1111/j.1600-0706.2011.20131.x

Chatfield, C., & Xing, H. (2019). The analysis of time series: An introduction with R (Seventh edition). CRC Press/Taylor and Francis Group.

Christie, M. R., Meirmans, P. G., Gaggiotti, O. E., Toonen, R. J., & White, C. (2017). Disentangling the relative merits and disadvantages of parentage analysis and assignment tests for inferring population connectivity. ICES Journal of Marine Science, 74(6), 1749–1762. 10.1093/icesjms/fsx044

Coelho, I., Castanheira, I., Bordado, J. M., Donard, O., & Silva, J. A. L. (2017). Recent developments and trends in the application of strontium and its isotopes in biological related fields. Trends in Analytical Chemistry, 90, 45–61. 10.1016/j.trac.2017.02.005

Cordoleani, F., Phillis, C. C., Sturrock, A. M., FitzGerald, A. M., Malkassian, A., Whitman, G. E., Weber, P. K., & Johnson, R. C. (2021). Threatened salmon rely on a rare life history strategy in a warming landscape. Nature Climate Change, 11(11), 982–988. 10.1038/s41558-021-01186-4

Courter, I. I., Child, D. B., Hobbs, J. A., Garrison, T. M., Glessner, J. J. G., & Duery, S. (2013). Resident rainbow trout produce anadromous offspring in a large interior watershed. Canadian Journal of Fisheries and Aquatic Sciences, 70(5), 701–710. 10.1139/cjfas-2012-0457

Cowen, R. K., & Sponaugle, S. (2009). Larval dispersal and marine population connectivity. Annual Review of Marine Science, 1(1), 443–466. 10.1146/annurev.marine.010908.163757

Dietterich, T. G. (2000). Ensemble methods in machine learning. In G. Goos, J. Hartmanis, & J. Van Leeuwen (Eds.), Multiple Classifier Systems (Vol. 1857, pp. 1–15). Springer Berlin Heidelberg. 10.1007/3-540-45014-9_1

Doubleday, Z. A., Martino, J. C., & Trueman, C. (2022). Harnessing universal chemical markers to trace the provenance of marine animals. Ecological Indicators, 144, 109481. 10.1016/j.ecolind.2022.109481

Elsdon, T. S., Wells, B. K., Campana, S. E., Gillanders, B. M., Jones, C. M., Limburg, K. E., Secor, D. H., Thorrold, S. R., & Walther, B. D. (2008). Otolith chemistry to describe movements and life-history parameters of fishes: Hypotheses, assumptions, limitations and inferences. Oceanography and Marine Biology: An Annual Review, 46, 297–330.

Gillanders, B. M. (2002). Temporal and spatial variability in elemental composition of otoliths: Implications for determining stock identity and connectivity of populations. Canadian Journal of Fisheries and Aquatic Sciences, 59(4), 669–679. 10.1139/f02-040

Giorgino, T. (2009). Computing and visualizing dynamic time warping alignments in *R*: The dtw package. Journal of Statistical Software, 31(7). 10.18637/jss.v031.i07

Gonzabato, N. (2025). manymodelr: Build and tune several models [Computer software]. https://CRAN.R-project.org/package=manymodelr

Hanson, N. N., Wurster, C. M., Eimf, & Todd, C. D. (2010). Comparison of secondary ion mass spectrometry and micromilling/continuous flow isotope ratio mass spectrometry techniques used to acquire intra-otolith *δ*^18^ O values of wild Atlantic salmon (*Salmo salar*). Rapid Communications in Mass Spectrometry, 24(17), 2491–2498. 10.1002/rcm.4646

Hegg, J. C., & Kennedy, B. P. (2021). Let’s do the time warp again: Non-linear time series matching as a tool for sequentially structured data in ecology. Ecosphere, 12(9). 10.1002/ecs2.3742

Hegg, J. C., Kennedy, B. P., & Chittaro, P. (2019). What did you say about my mother? The complexities of maternally derived chemical signatures in otoliths. Canadian Journal of Fisheries and Aquatic Sciences, 76(1), 81–94. 10.1139/cjfas-2017-0341

Hobson, K. A. (1999). Tracing origins and migration of wildlife using stable isotopes: A review. Oecologia, 120(3), 314–326. 10.1007/s004420050865

Huber, E. R., & Carlson, S. M. (2015). Temporal trends in hatchery releases of fall-run Chinook salmon in California’s Central Valley. San Francisco Estuary and Watershed Science, 13(2). 10.15447/sfews.2015v13iss2art3

Hüssy, K., Limburg, K. E., de Pontual, H., Thomas, O. R. B., Cook, P. K., Heimbrand, Y., Blass, M., & Sturrock, A. M. (2020). Trace element patterns in otoliths: The role of biomineralization. Reviews in Fisheries Science & Aquaculture. 10.1080/23308249.2020.1760204

Johnson, R. C., Weber, P. K., Wikert, J. D., Workman, M. L., MacFarlane, R. B., Grove, M. J., & Schmitt, A. K. (2012). Managed metapopulations: Do salmon hatchery ‘sources’ lead to in-river ‘sinks’ in conservation? PLoS ONE, 7(2), e28880. 10.1371/journal.pone.0028880

Johnson, R., Garza, J., MacFarlane, R., Grimes, C., Phillis, C., Koch, P., Weber, P., & Carr, M. (2016). Isotopes and genes reveal freshwater origins of Chinook salmon *Oncorhynchus tshawytscha* aggregations in California’s coastal ocean. Marine Ecology Progress Series, 548, 181–196. 10.3354/meps11623

Kuhn, M., & Wickham, H. (2020). Tidymodels*: A collection of packages for modeling and machine learning using tidyverse principles*. R package version 0.2.0.

Mercier, L., Darnaude, A. M., Bruguier, O., Vasconcelos, R. P., Cabral, H. N., Costa, M. J., Lara, M., Jones, D. L., & Mouillot, D. (2011). Selecting statistical models and variable combinations for optimal classification using otolith microchemistry. Ecological Applications, 21(4), 1352–1364. 10.1890/09-1887.1

Mokadem, F., Parkinson, I. J., Hathorne, E. C., Anand, P., Allen, J. T., & Burton, K. W. (2015). High-precision radiogenic strontium isotope measurements of the modern and glacial ocean: Limits on glacial–interglacial variations in continental weathering. Earth and Planetary Science Letters, 415, 111–120. 10.1016/j.epsl.2015.01.036

Neubauer, P., Shima, J. S., & Swearer, S. E. (2013). Inferring dispersal and migrations from incomplete geochemical baselines: Analysis of population structure using Bayesian infinite mixture models. Methods in Ecology and Evolution, 4(9), 836–845. 10.1111/2041-210X.12076

Newsome, S. D., Clementz, M. T., & Koch, P. L. (2010). Using stable isotope biogeochemistry to study marine mammal ecology. Marine Mammal Science. 10.1111/j.1748-7692.2009.00354.x

Pérez, G. E., & Hobson, K. A. (2007). Feather deuterium measurements reveal origins of migratory western loggerhead shrikes (*Lanius ludovicianus excubitorides*) wintering in Mexico. Diversity and Distributions, 13(2), 166–171. 10.1111/j.1472-4642.2006.00306.x

Phillis, C. C., Sturrock, A. M., Johnson, R. C., & Weber, P. K. (2018). Endangered winter-run Chinook salmon rely on diverse rearing habitats in a highly altered landscape. Biological Conservation, 217, 358–362. 10.1016/j.biocon.2017.10.023

R Core Team. (2025). R: a language and environment for statistical computing [Computer software]. https://www.R-project.org/

Reis-Santos, P., Gillanders, B. M., Sturrock, A. M., Izzo, C., Oxman, D. S., Lueders-Dumont, J. A., Hüssy, K., Tanner, S. E., Rogers, T., Doubleday, Z. A., Andrews, A. H., Trueman, C., Brophy, D., Thiem, J. D., Baumgartner, L. J., Willmes, M., Chung, M.-T., Charapata, P., Johnson, R. C., … Walther, B. D. (2023). Reading the biomineralized book of life: Expanding otolith biogeochemical research and applications for fisheries and ecosystem-based management. Reviews in Fish Biology and Fisheries, 33(2), 411–449. 10.1007/s11160-022-09720-z

Rosinski, C. L., Glaid, J., Hahn, M., & Fetzer, W. W. (2023). Natal origin differentiation using eye lens stable isotope analysis. North American Journal of Fisheries Management, 43(2), 547–555. 10.1002/nafm.10875

Schindler, D. E., Hilborn, R., Chasco, B., Boatright, C. P., Quinn, T. P., Rogers, L. A., & Webster, M. S. (2010). Population diversity and the portfolio effect in an exploited species. Nature, 465(7298), 609–612. 10.1038/nature09060

Secor, D. H. (1999). Specifying divergent migrations in the concept of stock: The contingent hypothesis. Fisheries Research, 43(1–3), 13–34. 10.1016/S0165-7836(99)00064-8

Secor, D. H. (2015). Migration ecology of marine fishes. Johns Hopkins university press.

Sturrock, A. M., Carlson, S. M., Wikert, J. D., Heyne, T., Nusslé, S., Merz, J. E., Sturrock, H. J. W., & Johnson, R. C. (2020). Unnatural selection of salmon life histories in a modified riverscape. Global Change Biology, 26(3), 1235–1247. 10.1111/gcb.14896

Sturrock, A. M., Hunter, E., Milton, J. A., EIMF, Johnson, R. C., Waring, C. P., & Trueman, C. N. (2015). Quantifying physiological influences on otolith microchemistry. Methods in Ecology and Evolution, 6(7), 806–816. 10.1111/2041-210X.12381

Sturrock, A. M., Satterthwaite, W. H., Cervantes-Yoshida, K. M., Huber, E. R., Sturrock, H. J. W., Nusslé, S., & Carlson, S. M. (2019). Eight decades of hatchery salmon releases in the California Central Valley: Factors influencing straying and resilience. Fisheries, 44(9), 433–444. 10.1002/fsh.10267

Sturrock, A. M., Wikert, J. D., Heyne, T., Mesick, C., Hubbard, A. E., Hinkelman, T. M., Weber, P. K., Whitman, G. E., Glessner, J. J., & Johnson, R. C. (2015). Reconstructing the migratory behavior and long-term survivorship of juvenile Chinook salmon under contrasting hydrologic regimes. PLOS ONE, 10(5), e0122380. 10.1371/journal.pone.0122380

Teichert, N., Tabouret, H., Lizé, A., Daverat, F., Acou, A., Trancart, T., Virag, L.-S., Pécheyran, C., Feunteun, E., & Carpentier, A. (2024). Quantifying larval dispersal portfolio in seabass nurseries using otolith chemical signatures. Marine Environmental Research, 196, 106426. 10.1016/j.marenvres.2024.106426

van Dijk, J. G. B., Meissner, W., & Klaassen, M. (2014). Improving provenance studies in migratory birds when using feather hydrogen stable isotopes. Journal of Avian Biology, 45(1), 103–108. 10.1111/j.1600-048X.2013.00232.x

Walther, B. D., & Limburg, K. E. (2012). The use of otolith chemistry to characterize diadromous migrations. Journal of Fish Biology, 81(2), 796–825. 10.1111/j.1095-8649.2012.03371.x

Walther, B. D., & Torrance, L. E. (2024). Quantifying euryhaline histories in red drum *Sciaenops ocellatus*: Otolith chemistry and muscle isotope ratios. Journal of Fish Biology, 105(5), 1389– 1405. 10.1111/jfb.15173

Wang, X., Mueen, A., Ding, H., Trajcevski, G., Scheuermann, P., & Keogh, E. (2013). Experimental comparison of representation methods and distance measures for time series data. Data Mining and Knowledge Discovery, 26(2), 275–309. 10.1007/s10618-012-0250-5

Williams, J. G. (2006). Central Valley salmon: A perspective on Chinook and steelhead in the Central Valley of California. San Francisco Estuary and Watershed Science, 4(3). 10.15447/sfews.2006v4iss3art2

Willmes, M., Sturrock, A. M., Cordoleani, F., Hugentobler, S., Meek, M. H., Whitman, G., Evans, K., Palkovacs, E. P., Stauffer-Olsen, N. J., & Johnson, R. C. (2024). Integrating otolith and genetic tools to reveal intraspecific biodiversity in a highly impacted salmon population. Journal of Fish Biology, 105(2), 412–430. 10.1111/jfb.15847

Wright, M. N., & Ziegler, A. (2017). ranger: A fast implementation of random forests for high dimensional data in C++ and R. Journal of Statistical Software, 77(1). 10.18637/jss.v077.i01

