## Supplementary Material for "Advancing Provenance Assignment using Machine Learning and Time Series Analysis of Chemical Chronologies in Archival Tissues"

### **Supplementary Text**

##### Text S1. Synthetic early migrant simulation

The current reference library of known-origin Central Valley salmon otoliths lacks samples of “early migrants” – fish that emigrated soon after emergence as fry, then reared in non-natal habitats. These early migrants are typically too small to tag, and thus their natal origin cannot be determined if sampled downstream. However, excluding this phenotype from the training dataset can introduce substantial bias given that they often dominate juveniles leaving the natal stream (Sturrock et al., 2020). While early migrants can potentially rear in any habitat downstream of their natal river in the Central Valley, they generally leave the natal stream during high winter flows and rear in the lower reaches of the mainstem Sacramento and San Joaquin Rivers within the freshwater Delta (Sturrock et al., 2020; Williams, 2006). To account for this, we conducted a simulation to generate synthetic otolith ^87^Sr/^86^Sr profiles for juvenile Chinook salmon early migrants, using established biological principles related to fish movement patterns and known water chemistries.

The model was informed by known water ^87^Sr/^86^Sr ranges for different natal rivers, as well as the size and timing of juvenile salmon movements derived from existing data for fall-run and spring-run Chinook salmon (Table S2, which summarizes the input parameters to generate synthetic early migrants). For each fish, we generated a high-resolution ^87^Sr/^86^Sr profile from 0 µm (the otolith core) to 500 µm, at 2.5 µm intervals. We then binned these data into 40 µm intervals by averaging the values within each bin. This binning process reduced variance in the ^87^Sr/^86^Sr profile and aligned the profile with the temporal resolution of our empirical otolith laser-ablation data. These profiles included distinct stages of juvenile otolith formation, including the core region, emergence, natal river rearing, and downstream rearing (Fig. 1, Table S2).

For the otolith core region, we assumed an initial marine ^87^Sr/^86^Sr value typical of fall-run Chinook salmon, reflecting maternal yolk deposition at sea, transitioning linearly to the natal river value by the time of emergence. For sources where spring-run fish were used as reference data (BUT, MIL, and DEE), ^87^Sr/^86^Sr values were drawn from simulated normal distributions based on the mean and standard deviation of otolith core ^87^Sr/^86^Sr measurements of known-origin spring-run samples collected from each source. This approach reflects the tendency of spring-run offspring to exhibit core values between marine and natal signatures, due to their parents’ prolonged freshwater residence prior to spawning (Williams, 2006). From emergence to outmigration, otolith profiles were assigned ^87^Sr/^86^Sr values drawn from a simulated normal distribution based on natal river water samples. Early migrants were modeled to outmigrate after reaching an otolith size of approximately 250 ± 50 μm (range: 175–330 μm), after which they were assigned ^87^Sr/^86^Sr values representative of downstream rearing habitats, also drawn from distributions based on water chemistry data. The total otolith growth during downstream rearing was modeled as a normal distribution with a mean of 250 μm and a standard deviation of 40 μm.

To ensure our training data represented a broad range of life histories, including early migrants, we generated synthetic ^87^Sr/^86^Sr profiles for all natural sources. Hatcheries were excluded from this process, as no Central Valley hatchery currently releases salmon smaller than 60 mm (Huber & Carlson, 2015). Thus, we generated and added five synthetic early migrant profiles per natural source to the training dataset (Table S1), bringing the total to 15 samples per natural source (10 empirical otolith + 5 synthetic data). Hatchery sources included 15 empirical otolith samples each.

##### Text S2. Otolith microstructure analysis

Due to differences in food availability for newly emerged salmon in rivers versus hatcheries, otoliths from natural-origin fish often exhibit a distinct “stress” check at the point of yolk sac depletion—a feature typically absent in hatchery-reared fish. This characteristic has been previously used to estimate hatchery versus natural contributions to the ocean fishery (Barnett-Johnson et al., 2007) and was used here as an additional predictor in the assignment model.

All otoliths in the reference library were visually inspected under transmitted light at 5× or 10× magnification. The presence of the exogenous feeding check was scored independently by at least two readers using a three-point scale: a score of 2 indicated a distinct, sharp stress check occurring approximately 200–250 µm from the core; a score of 1 indicated the absence of a check or the presence of an indistinct check at this distance; and a score of 1.5 was assigned when the sample was unusable or the check was unclear. The scores were then averaged to produce a mean exogenous feeding score for each otolith. Synthetic early migrants from natural-origin sources were assigned the mean score calculated across all natural-origin fish. To assess the contribution of this variable in model performance, all assignment models were also trained and evaluated without the mean exogenous feeding score as a predictor.

The addition of otolith microstructure features as a predictor alongside otolith ^87^Sr/^86^Sr in the natal assignment model only marginally improved model performance, except in the automated median approach, where it provided a notable increase (Fig. S5). Thus, this additional step may be omitted in future applications to maximize efficiency, particularly when using models that incorporate time series data.

##### Text S3. Comparison of machine learning models with traditional statistical classifiers

To evaluate the performance of machine learning models such as Random Forest (RF) relative to traditional statistical classifiers, we compared classification accuracy for the automated median approach (Fig. 3C) implemented using RF, linear discriminant analysis (LDA), and quadratic discriminant analysis (QDA). This comparison was restricted to the automated median approach, as LDA and QDA could not be applied to models with larger sets of predictors due to collinearity and rank deficiency, which limited their use to this simplified configuration.

Using 10-fold cross-validation, we found that the classification accuracy of the automated median approach was comparable between RF (accuracy = 0.409) and QDA (0.407), whereas LDA performed substantially worse (accuracy = 0.283; Fig. S6).

##### Text S4. User-defined natal region comparison

To evaluate the subjectivity involved in user-defined natal region assignments—and how these factors might influence subsequent natal assignment results—we conducted a natal region identification trial with three independent readers. All readers were trained before the exercise and were experienced in visually inspecting otolith ^87^Sr^/86^Sr profiles of salmonids. Each reader was tasked with identifying a natal region for each fish by visually inspecting its otolith ^87^Sr^/86^Sr profile, without any prior knowledge of the fish (i.e., collection site, collection year, fish FL).

Using known-origin juvenile samples (n = 255), we compared the starting otolith distance of the assigned natal region and the median ^87^Sr^/86^Sr value within the user-defined natal region across the three readers. Paired *t*-tests were conducted to assess differences in matched pairs of values among the three readers. Additionally, through 10-fold cross-validation (repeated 10 times), we compared the classification accuracy of the “user-defined median approach” using the median ^87^Sr/^86^Sr values from the three readers. Pairwise classification agreement was used to assess consistency in natal assignments among the readers.

The natal region identification trial and paired *t*-tests indicated that the starting distance of the natal region in the otolith ^87^Sr/^86^Sr profile differed significantly among the three readers (Table S4A; Fig. S7A). Specifically, while readers 1 and 3 generally aligned with the identity line in the agreement plot, reader 2 showed a systematic bias, assigning the natal region at a later point in the otolith ^87^Sr/^86^Sr profile.

The difference in the starting distance of the natal region did not necessarily result in differences in the median ^87^Sr/^86^Sr values within the user-defined natal region (Table S4B; Fig. S7B). No significant differences were detected in the matched pairs of median ^87^Sr/^86^Sr values among the three readers, despite reader 2’s tendency to assign the natal region at a later point in the otolith ^87^Sr/^86^Sr profile.

The cross-validated mean classification accuracy among the three readers using the “user-defined median approach” ranged from 0.458 to 0.476 (Table S4C). All readers achieved similar precision (1 SD) ranging from 0.013 to 0.018. The overall natal assignment agreement ranged from 0.582 to 0.666, indicating that the assignment results were largely inconsistent and strongly influenced by each reader’s decision on natal region identification (Table S4D).

### **Supplementary Tables and Figures**

##### **Table S1.** Summary of otolith ^87^Sr/^86^Sr profile datasets used for model training and application. Each row represents a unique combination of site, origin (natural or hatchery), and sample type. The “Objective” column indicates whether (i) samples were used to train the classification model, (ii) apply the model to unknown-origin juveniles, or (iii) evaluate the impact of including synthetic early migrant data by testing model performance on known-origin adult returners. Collection year indicates the range of years in which samples were collected.

| Objective | Site Name | Site Code | Natural or Hatchery | Sample Type | n | Collection Year |
| --- | --- | --- | --- | --- | --- | --- |
| Model training | Sacramento River | SAC | Natural | Adult winter run (assumed return) | 10 | 2006-2015 |
|  | Sacramento River | SAC | Natural | Synthetic early migrant | 5 | Not applicable |
|  | Coleman National Fish Hatchery | CNH | Hatchery | CWT tagged adult hatchery fish | 2 | 2009-2010 |
|  | Coleman National Fish Hatchery | CNH | Hatchery | CWT tagged juvenile hatchery fish sample downstream of source | 6 | 2014-2017 |
|  | Coleman National Fish Hatchery | CNH | Hatchery | Juvenile sampled at source before outmigration | 7 | 2000-2015 |
|  | Battle Creek | BAT | Natural | Juvenile sampled at source before outmigration | 10 | 1999-2014 |
|  | Battle Creek | BAT | Natural | Synthetic early migrant | 5 | Not applicable |
|  | Mill Creek | MIL | Natural | Adult spring run (assumed return) | 10 | 2010-2011 |
|  | Mill Creek | MIL | Natural | Synthetic early migrant | 5 | Not applicable |
|  | Deer Creek | DEE | Natural | Adult spring run (assumed return) | 10 | 2008-2018 |
|  | Deer Creek | DEE | Natural | Synthetic early migrant | 5 | Not applicable |
|  | Butte Creek | BUT | Natural | Adult spring run (assumed return) | 1 | 2018 |
|  | Butte Creek | BUT | Natural | Juvenile sampled at source before outmigration | 9 | 2018 |
|  | Butte Creek | BUT | Natural | Synthetic early migrant | 5 | Not applicable |
|  | Feather River | FEA | Natural | Juvenile sampled at source before outmigration | 10 | 1999-2019 |
|  | Feather River | FEA | Natural | Synthetic early migrant | 5 | Not applicable |
|  | Feather River Hatchery | FEH | Hatchery | CWT tagged adult hatchery fish | 7 | 2007-2008 |
|  | Feather River Hatchery | FEH | Hatchery | CWT tagged juvenile hatchery fish sample downstream of source | 4 | 2014-2018 |
|  | Feather River Hatchery | FEH | Hatchery | Juvenile sampled at source before outmigration | 4 | 2002-2019 |
|  | Yuba River | YUB | Natural | Juvenile sampled at source before outmigration | 10 | 2002-2019 |
|  | Yuba River | YUB | Natural | Synthetic early migrant | 5 | Not applicable |
|  | American River | AME | Natural | Juvenile sampled at source before outmigration | 10 | 1999-2018 |
|  | American River | AME | Natural | Synthetic early migrant | 5 | Not applicable |
|  | Nimbus Fish Hatchery | NIH | Hatchery | CWT tagged adult hatchery fish | 1 | 2017 |
|  | Nimbus Fish Hatchery | NIH | Hatchery | CWT tagged juvenile hatchery fish sample downstream of source | 4 | 2016-2018 |
|  | Nimbus Fish Hatchery | NIH | Hatchery | Juvenile sampled at source before outmigration | 10 | 2002-2019 |
|  | Mokelumne River | MOK | Natural | Juvenile sampled at source before outmigration | 10 | 2000-2017 |
|  | Mokelumne River | MOK | Natural | Synthetic early migrant | 5 | Not applicable |
|  | Mokelumne River Hatchery | MOH | Hatchery | CWT tagged adult hatchery fish | 7 | 2001-2017 |
|  | Mokelumne River Hatchery | MOH | Hatchery | CWT tagged juvenile hatchery fish sample downstream of source | 1 | 2018 |
|  | Mokelumne River Hatchery | MOH | Hatchery | Juvenile sampled at source before outmigration | 7 | 1999-2017 |
|  | Stanislaus River | STA | Natural | Juvenile sampled at source before outmigration | 10 | 1999-2017 |
|  | Stanislaus River | STA | Natural | Synthetic early migrant | 5 | Not applicable |
|  | Tuolumne River | TUO | Natural | Juvenile sampled at source before outmigration | 10 | 2003-2018 |
|  | Tuolumne River | TUO | Natural | Synthetic early migrant | 5 | Not applicable |
|  | Merced River | MER | Natural | Juvenile sampled at source before outmigration | 10 | 2003-2014 |
|  | Merced River | MER | Natural | Synthetic early migrant | 5 | Not applicable |
|  | Merced River Hatchery | MEH | Hatchery | CWT tagged adult hatchery fish | 3 | 2003-2009 |
|  | Merced River Hatchery | MEH | Hatchery | CWT tagged juvenile hatchery fish sample downstream of source | 6 | 2015-2016 |
|  | Merced River Hatchery | MEH | Hatchery | Juvenile sampled at source before outmigration | 6 | 2002-2017 |
| Model application - unknown-origin juvenile | Chipps Island | NA | NA | Unknown origin juvenile | 50 | 2014-2021 |
| Model application - known-origin adult returners | Yuba River | YUB | Natural | Known-origin adult returners that outmigrated early as juveniles | 10 | 2009-2019 |
|  | American River | AME | Natural | Known-origin adult returners that outmigrated early as juveniles | 10 | 2014-2021 |
|  | Stanislaus River | STA | Natural | Known-origin adult returners that outmigrated early as juveniles | 10 | 2001-2013 |

##### **Table S2.** Parameters used to generate synthetic early migrants to train the assignment model, with all distances representing the radius of the sagittal otolith from the primordia to the dorsal edge.

| **Parameter** | **Value** | **Justification/source** |
| --- | --- | --- |
| Profile length | 500 µm | The distance was chosen to encompass all major stages of the early migrant phenotype of juvenile Chinook salmon otolith development, including the core region, emergence, natal river rearing, and early downstream rearing. |
| Step size | 40 µm | Matches temporal resolution of empirical otolith laser ablation data. |
| Core size | 20 µm | This was calculated as half the laser spot diameter (40 µm), reflecting the typical starting point of laser-ablation profiles at the otolith core. |
| Core to emergence | Linear interpolation | This assumption is based on the observed isotopic shift from the core (reflecting maternal marine input) to the natal river signature at emergence. The actual rate of this transition (slope) depends on the ^87^Sr/^86^Sr values in both the otolith core and the natal river water. |
| Otolith size at emergence | Normal distribution (mean 225 ± 30 µm SD) | Midpoint between two independent datasets with exogenous feeding check distance estimated (mean ± SD = 217.1 ± 24.5 µm, n = 3113, and 236.1 ± 28.0 µm, n = 960; Sturrock et al., unpublished) |
| Otolith size for natal exit size of early migrants | Normal distribution (mean 250 ± 50 µm SD, lower limit = 175 µm, upper limit = 330 µm) | Early migrants typically range from 30–45 mm in length at the time of natal exit (Williams, 2006 and references therin), which corresponds to an otolith radius of approximately 200–300 µm, based on equations from Sturrock et al., (2020) and Willmes et al., (2024). The smallest early migrants observed in various unpublished datasets had otolith radii as small as 180 µm, thus we set the lower limit at 175 µm. |
| Total otolith growth during downstream rearing | Normal distribution (mean 250 ± 40 µm SD) | Based on the mean otolith radii at natal and freshwater exit observed by Cordoleani et al., (2021) and Willmes et al., (2024). |
| Fall-run core ^87^Sr/^86^Sr value | Normal distribution (mean 0.70918 ± 0.0001 SD) | Marine value from Mokadem et al., 2015). |
| Spring-run core ^87^Sr/^86^Sr value | Normal distribution BUT (mean 0.70722 ± 0.0004 SD), DEE (mean 0.70650 ± 0.0006 SD), MIL (mean 0.70650 ± 0.0004 SD) | Based on the mean and standard deviation of otolith core ^87^Sr/^86^Sr measurements of known-origin spring-run samples collected from Butte (BUT), Deer (DEE), and Mill (MIL) Creeks. |
| Natal river ^87^Sr/^86^Sr value | Normal distribution (^87^Sr/^86^Sr mean ± SD) | Local natal river ^87^Sr/^86^Sr value reported in part by Barnett-Johnson et al., (2008); Phillis et al., (2018); Sturrock et al., (2020); Sturrock et al., (2015). |
| Downstream rearing ^87^Sr/^86^Sr value | Normal distribution (mean 0.70600 ± 0.0004 SD) | Local downstream ^87^Sr/^86^Sr values reported in part by Barnett-Johnson et al., (2008); Phillis et al., (2018); Sturrock et al., (2020); Sturrock et al., (2015). We assumed that early migrants moved directly to the central, freshwater region of the Sacramento–San Joaquin River Delta. |

##### **Table S3**. Optimal hyperparameters selected for natal assignment models. The *number of trees*, *minimum node size*, and *mtry* were tuned for the random forest models, while the number of nearest neighbors (*k*) was tuned for the dynamic time warping combined with *k*-nearest neighbors (DTW + KNN) approach.

| Assignment Method | Number of Trees | Minimum Node Size | mtry | k |
| --- | --- | --- | --- | --- |
| Automated Median | 1000 | 1 | 2 | NA |
| User-defined Median | 1000 | 20 | 2 | NA |
| Automated Feature | 1000 | 2 | 3 | NA |
| User-defined Feature | 1000 | 1 | 3 | NA |
| Discrete Time series | 1000 | 1 | 5 | NA |
| DTW + KNN | NA | NA | NA | 5 |

##### **Table S4.** Summary of user-defined natal region comparisons. This table includes results from paired *t*-tests comparing (A) starting distances of the natal region in the otolith ^87^Sr/^86^Sr profile, (B) median ^87^Sr/^86^Sr values, (C) cross-validated classification accuracy, and (D) natal assignment agreement among different readers.

| A. **Natal Region Starting Distance (µm)** | Reader | Statistic | Degrees of Freedom | *p*-value | Significance |
| --- | --- | --- | --- | --- | --- |
|  | R1-R2 | −11.1 | 254 | <0.001 | *** |
|  | R1-R3 | 4.23 | 254 | <0.001 | *** |
|  | R2-R3 | 11.4 | 254 | <0.001 | *** |
| B. **Median ^87^Sr/^86^Sr** | Reader | Statistic | Degrees of Freedom | *p*-value |  |
|  | R1-R2 | −0.09 | 254 | 1 | ns |
|  | R1-R3 | 0.706 | 254 | 1 | ns |
|  | R2-R3 | 0.619 | 254 | 1 | ns |
| C. **Classification Accuracy** | Reader | Mean | Standard Deviation |  |  |
|  | R1 | 0.476 | 0.018 |  |  |
|  | R2 | 0.458 | 0.017 |  |  |
|  | R3 | 0.482 | 0.013 |  |  |
| D. **Natal Assignment Agreement** | Reader | % Agreement |  |  |  |
|  | R1-R2 | 0.656 |  |  |  |
|  | R1-R3 | 0.666 |  |  |  |
|  | R2-R3 | 0.582 |  |  |  |

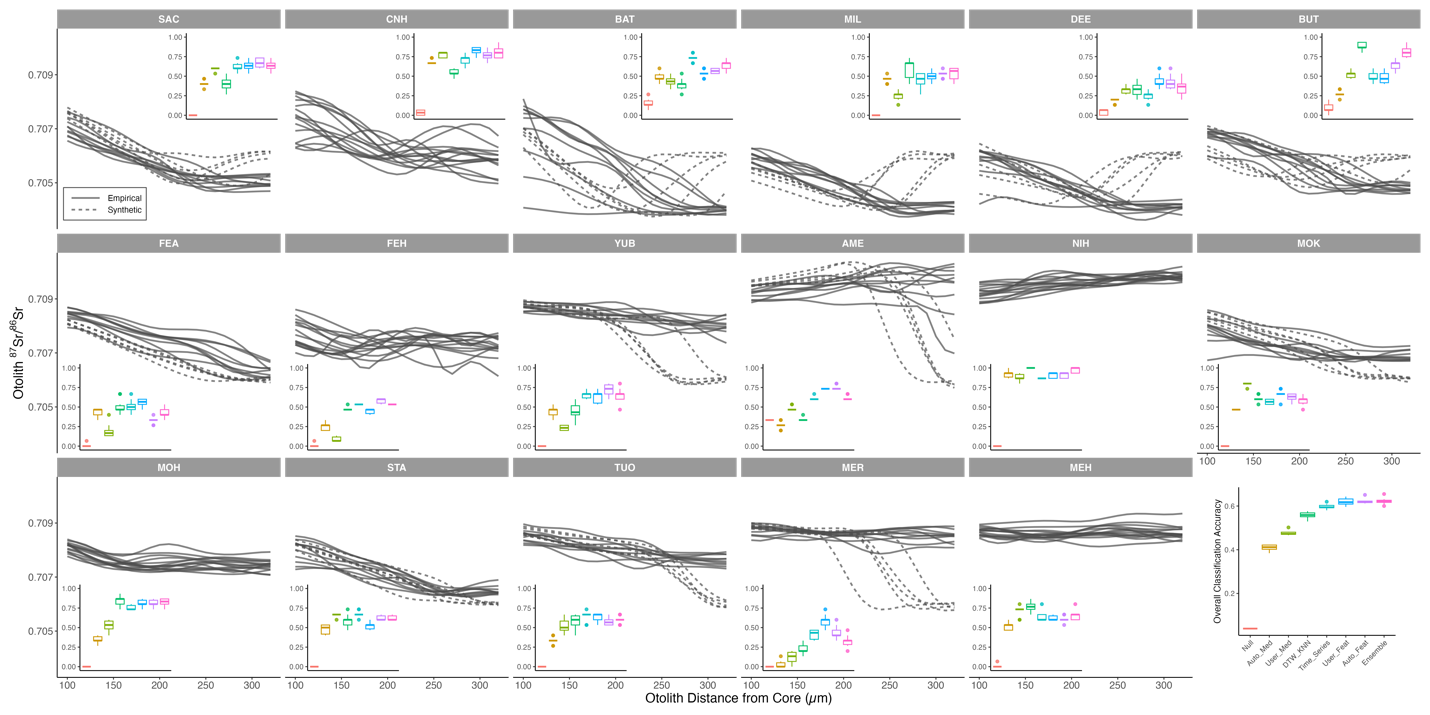

##### **Figure S1.** Classification accuracy of natal assignment models evaluated using repeated 10-fold cross-validation (n = 255 known-origin Chinook salmon). Overall classification accuracy is displayed in the bottom right corner. Inset panels show otolith ^87^Sr/^86^Sr profiles by natal source with accuracy for each. Solid and dashed lines represent otolith ^87^Sr/^86^Sr values for “empirical” and “synthetic” fish, respectively. Natal source and assignment model abbreviations are defined in Fig. 2 and Fig. 4, respectively.

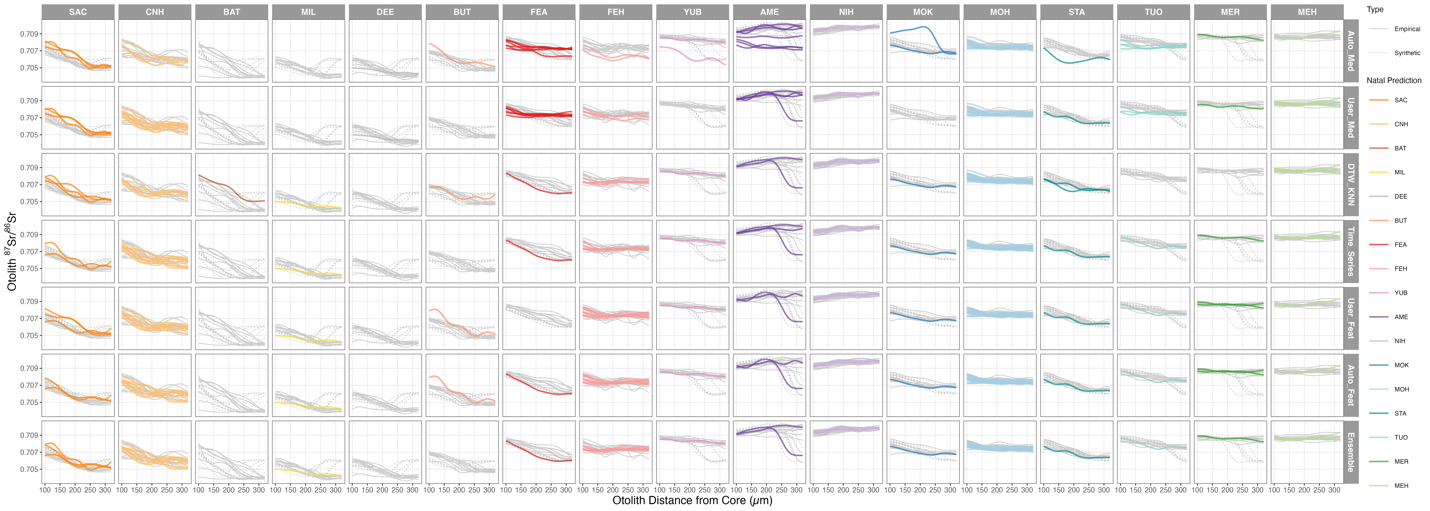

##### **Figure S2.** Otolith ^87^Sr/^86^Sr profiles and natal assignment results for unknown-origin juveniles collected at Chipps Island (n = 50) across different natal assignment models. Background gray lines represent the training data (solid for empirical and dashed for empirical samples), while colored lines indicate unknown-origin fish with their predicted natal origins. Natal source and assignment model abbreviations are defined in Fig. 2 and Fig. 4, respectively.

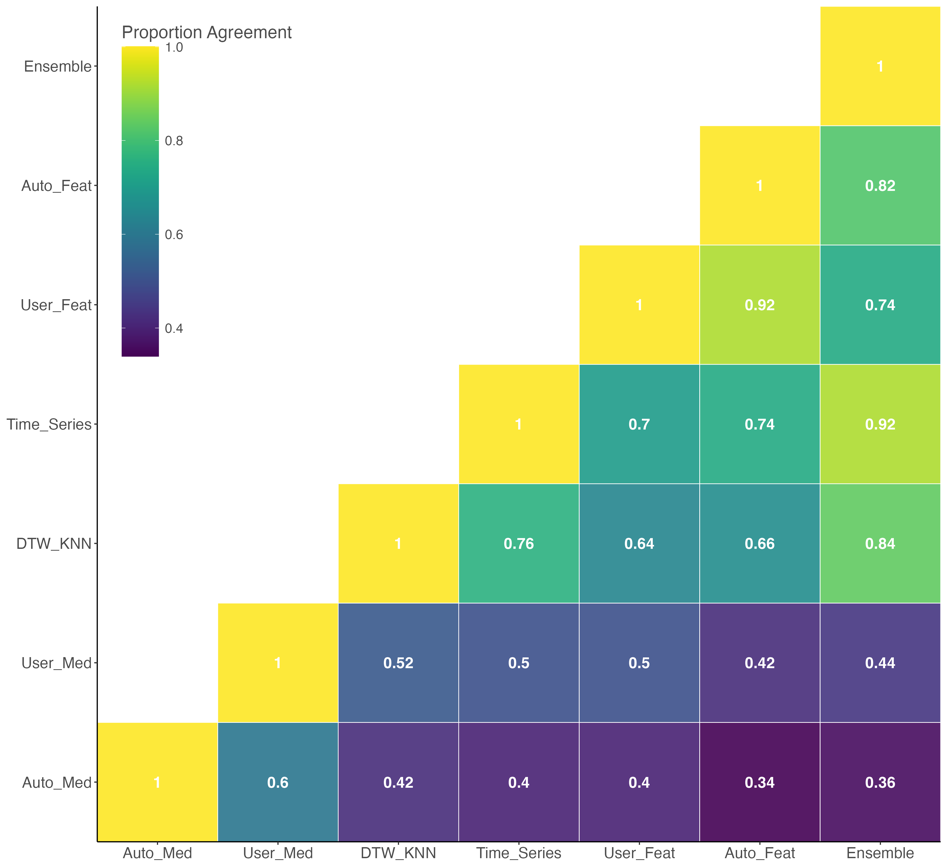

##### **Figure S3.** Natal assignment agreement matrix for pairs of different natal assignment models for unknown-origin juvenile samples collected at Chipps Island. Values within each grid indicate the proportion of agreement between different models. Assignment model abbreviations are defined in Fig. 4.

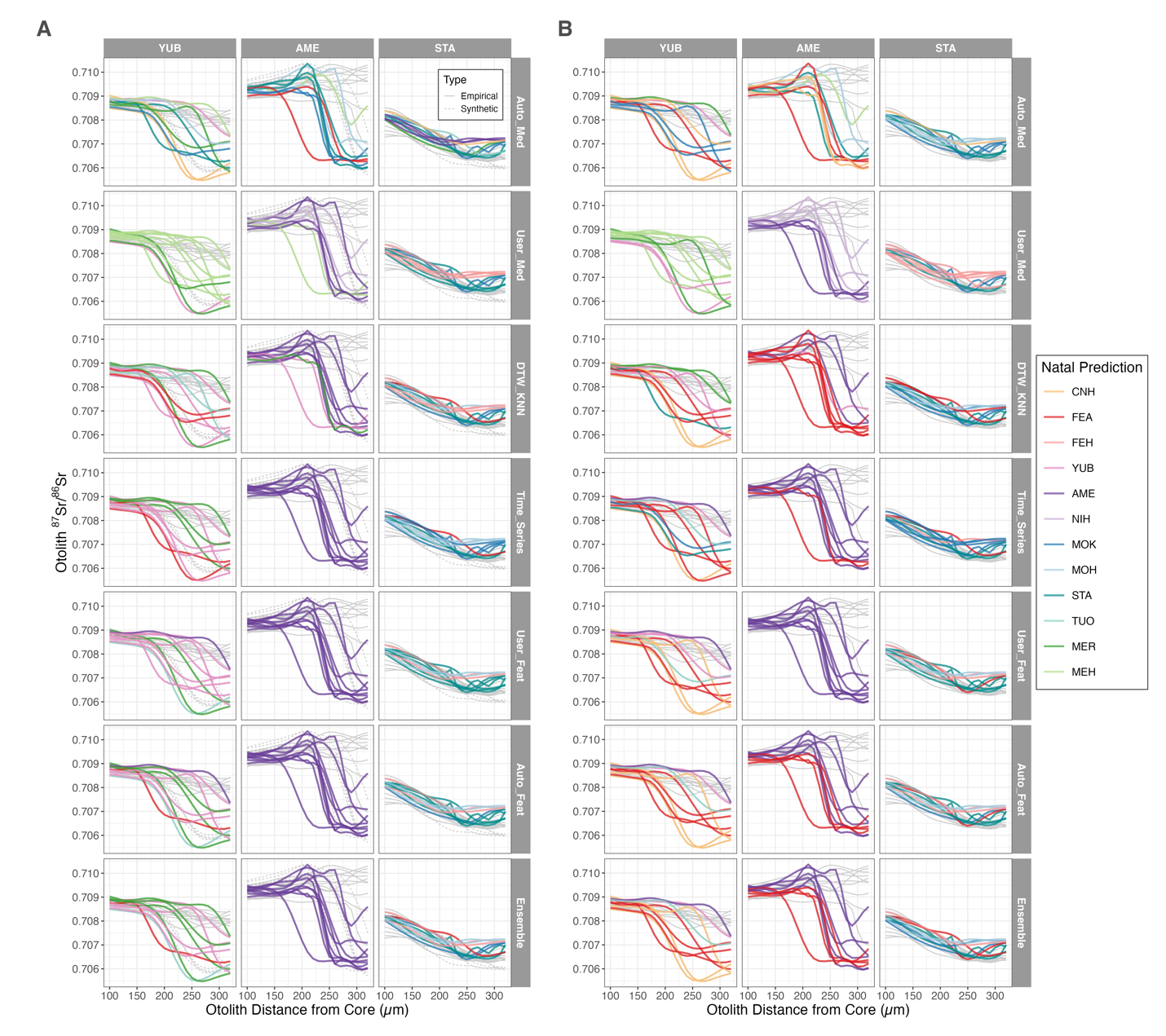

##### **Figure S4.** Otolith ^87^Sr/^86^Sr profiles and natal origin predictions for known-origin adult returns from the Yuba (YUB), American (AME), and Stanislaus (STA) Rivers and training dataset with (A) and without (B) synthetic early migrants. Panels are faceted by the true natal origin of each fish and colored by their predicted natal origin. Gray lines in the background represent the training data, with solid lines indicating empirical samples and dashed lines indicating synthetic early migrants. Natal source and assignment model abbreviations are defined in Fig. 2 and Fig. 4, respectively.

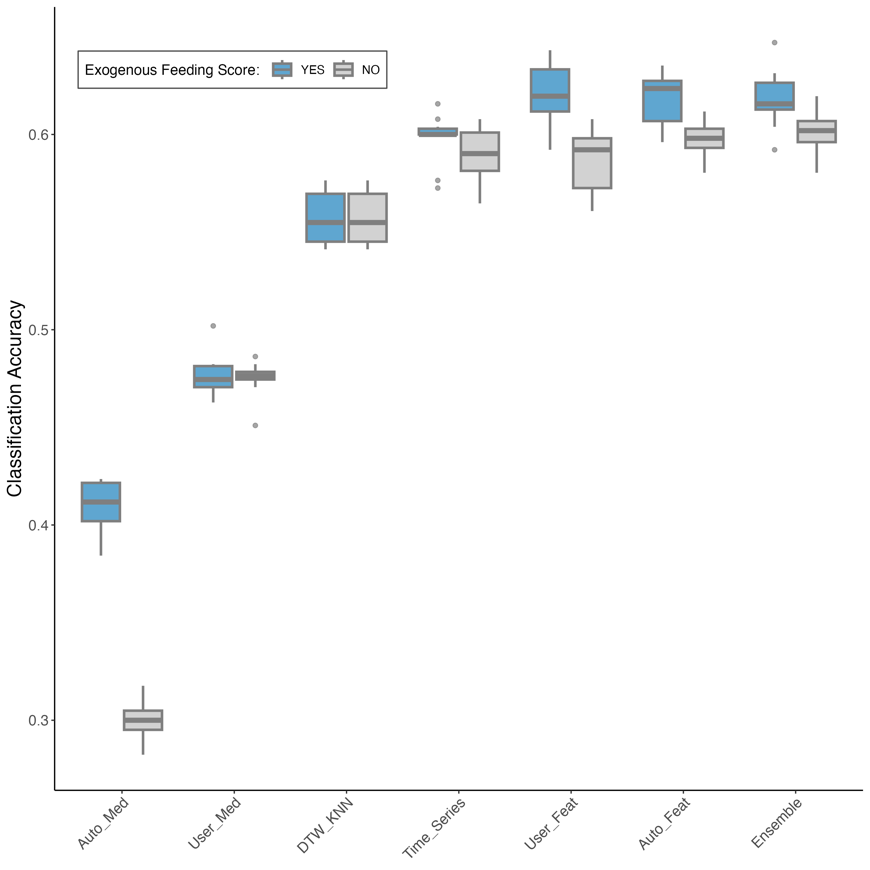

##### **Figure S5.** Boxplots showing classification accuracy for different natal assignment models trained with and without the otolith microstructure feature (exogenous feeding check score) as a predictor. Note that the DTW + KNN approach did not use this feature in either case, resulting in identical performance across both models. Assignment model abbreviations are defined in Fig. 4.

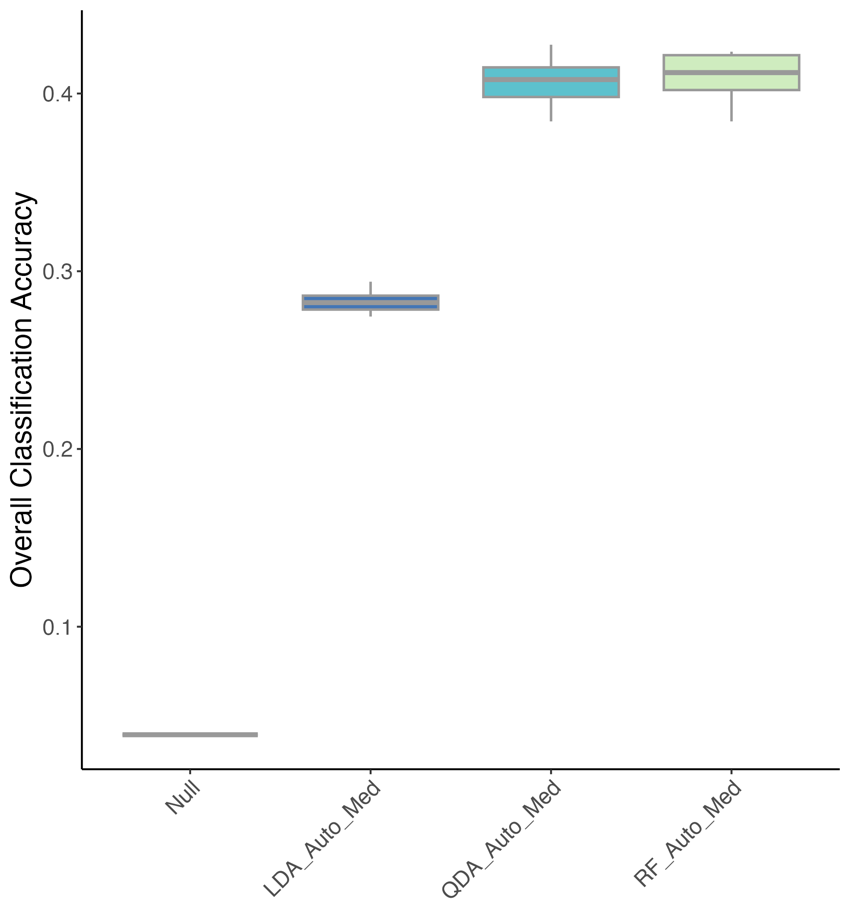

##### **Figure S6.** Boxplot showing overall classification accuracy for the automated median approach implemented using various classification methods, including linear discriminant analysis (LDA), quadratic discriminant analysis (QDA), and random forest (RF), evaluated using 10-fold cross-validation repeated 10 times on known-origin Chinook salmon (n = 255). Natal assignment model abbreviations: “Null” = non-informative null model; “LDA_Auto_Med” = automated median approach using LDA; “QDA_Auto_Med” = automated median approach using QDA; “RF_Auto_Med” = automated median approach using RF.

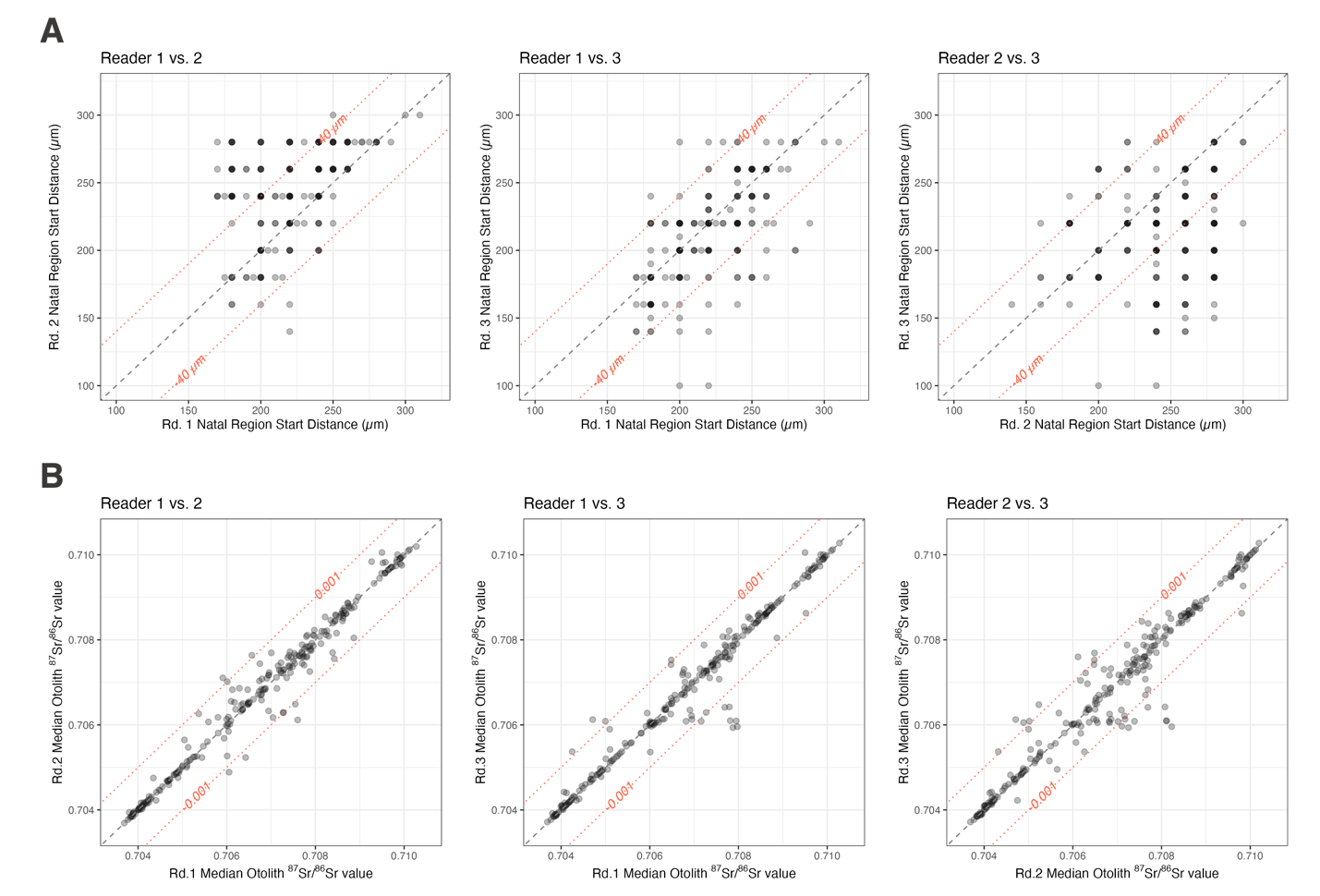

##### **Figure S7**. Agreement plots illustrating matched pairs of (A) starting distances of the natal region in the otolith ^87^Sr/^86^Sr profile and (B) median ^87^Sr/^86^Sr values within the user-defined natal region among three readers. The identity line is shown in gray dashed lines, with the offset of 40 µm for starting distances and 0.001 for median ^87^Sr/^86^Sr values indicated by red dotted lines.
